# Molecular identification of sixty-three amacrine cell types completes a mouse retinal cell atlas

**DOI:** 10.1101/2020.03.10.985770

**Authors:** Wenjun Yan, Mallory A. Laboulaye, Nicholas M. Tran, Irene E. Whitney, Inbal Benhar, Joshua R. Sanes

## Abstract

Amacrine cells (ACs) are a diverse class of interneurons that modulate input from photoreceptors to retinal ganglion cells (RGCs), rendering each RGC type selectively sensitive to particular visual features, which are then relayed to the brain. While many AC types have been identified morphologically and physiologically, they have not been comprehensively classified or molecularly characterized. We used high-throughput single-cell RNA sequencing (scRNA-seq) to profile >32,000 ACs from mouse retina, and applied computational methods to identify 63 AC types. We identified molecular markers for each type, and used them to characterize the morphology of multiple types. We show that they include nearly all previously known AC types as well as many that had not been described. Consistent with previous studies, most of the AC types express markers for the canonical inhibitory neurotransmitters GABA or glycine, but several express neither or both. In addition, many express one or more neuropeptides, and two express glutamatergic markers. We also explored transcriptomic relationships among AC types and identified transcription factors expressed by individual or multiple closely related types. Noteworthy among these were *Meis2* and *Tcf4*, expressed by most GABAergic and most glycinergic types, respectively. Together, these results provide a foundation for developmental and functional studies of ACs, as well as means for genetically accessing them. Along with previous molecular, physiological and morphological analyses, they establish the existence of at least 130 neuronal types and nearly 140 cell types in mouse retina.

**SIGNIFICANCE STATEMENT:** The mouse retina is a leading model for analyzing the development, structure, function and pathology of neural circuits. A complete molecular atlas of retinal cell types provides an important foundation for these studies. We used high-throughput single-cell RNA sequencing (scRNA-seq) to characterize the most heterogeneous class of retinal interneurons, amacrine cells, identifying 63 distinct types. The atlas includes types identified previously as well as many novel types. We provide evidence for use of multiple neurotransmitters and neuropeptides and identify transcription factors expressed by groups of closely related types. Combining these results with those obtained previously, we proposed that the mouse retina contains 130 neuronal types, and is therefore comparable in complexity to other regions of the brain.

## INTRODUCTION

The mouse retina has emerged as a leading model for studying the development, structure and function of neural circuits in the vertebrate central nervous system (Sanes and Masland, 2015; Seabrook et al., 2017). In addition, it is often used to investigate mechanisms underlying retinal disease, the major cause of irreversible blindness. An atlas of mouse retinal cell types would be a valuable resource for pursuing such studies. High-throughput single-cell RNA sequencing (scRNA-seq) is a promising method for achieving this goal: it enables comprehensive identification and molecular characterization of the cell types that comprise complex tissues, as well as a framework for incorporating structural and physiological data required for generating a definitive atlas (Zeng and Sanes, 2017). Moreover, it provides molecular markers that facilitate development of genetic strategies to access and manipulate specific cell types within neural circuits.

In an initial study, we used scRNA-seq to profile 44,808 cells from mouse retina, recovering the 6 major classes of cells present in vertebrate retinas: photoreceptors (PRs) that sense light; three classes of interneurons (horizontal cells, bipolar cells and amacrine cells – HCs, BCs and ACs) that receive information from photoreceptors and process it; retinal ganglion cells (RGCs) that receive information from interneurons and transmit it to central targets; and Müller glial cells (Macosko et al., 2015) (Figure 1A). This study was unable, however, to resolve all of the cell types into which the classes are divided: only 33 neuronal groups were recovered, even though the number of authentic types had been estimated to exceed 60 (Masland, 2012). The reason was that ∼80% of retinal cells are rod photoreceptors (Jeon et al., 1998), leaving too few of the less abundant but more diverse neuronal classes (BCs, ACs, and RGCs) to recover rare types or distinguish similar types from each other. Accordingly, we set out to enrich BCs, RGCs and ACs so we could profile them in sufficient numbers. For BCs and RGCs, we documented the existence of 15 and 46 types, respectively (Shekhar et al., 2016; Tran et al., 2019). These numbers correspond well to those obtained from recent high-throughput electrophysiological, ultrastructural and molecular studies (Baden et al., 2016; Bae et al., 2018; Franke et al., 2017; Greene et al., 2016; Rheaume et al., 2018).

**Figure 1.**
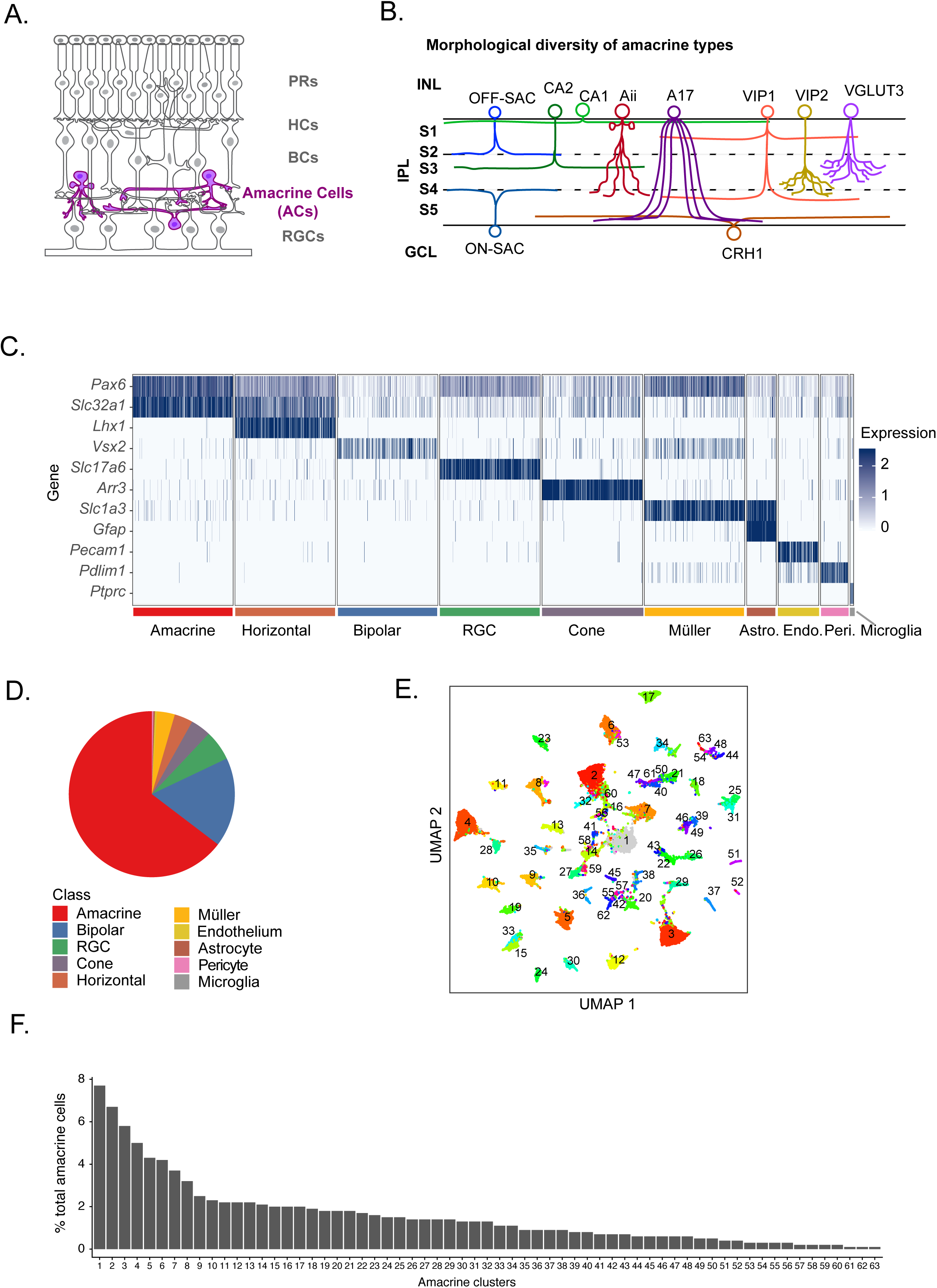
Single cell transcriptomes of mouse amacrine cells. A. Sketch of a retinal section showing cell classes and layers (Adapted from Tran et al. 2019). PR, photoreceptor; HC, horizontal cell; BC, bipolar cell; AC, amacrine cell; RGC, retinal ganglion cell. B. Sketch of the dendritic lamination patterns for various previously defined AC types. INL, inner nuclear layer; IPL, inner plexiform layer; GCL, ganglion cell layer. IPL sublaminae represented as S1-5. C. Expression patterns of marker genes used to allocate retinal cells to classes. Plot shows a randomly down-sampled subset of all cells. Color bars indicate cell class. D. Fraction of cells in each cell class, as determined by expression of canonical markers in C. E. Uniform Manifold Approximation and Projection (UMAP) visualization of 63 AC clusters. F. Relative frequencies of AC clusters, expressed as percentage of all 32,523 AC cells profiled. Clusters are numbered in order of decreasing frequency.

Here, we present an analysis of ACs. ACs receive synaptic input from BCs and other ACs, and provide output to BCs, other ACs and RGCs. They modify the visual signals that travel from photoreceptors to RGCs via BCs, thereby shaping the visual features to which each RGC type responds. Several AC types have been shown to play specific roles in retinal computation; for example, some render RGCs selectively responsive to motion in particular directions, and others able to distinguish local from global motion (Werblin., 2010; Vaney et al., 2012; Krishnaswamy et al., 2014; Lee et al., 2016; Tian et al., 2016; Diamond, 2017; Wei, 2018). Such roles require multiple AC types; indeed, they are generally thought to be the most heterogeneous retinal class (MacNeil and Masland, 1998; Lin and Masland, 2006) (Figure 1B).

Our transcriptomic analysis revealed 63 AC types, enabling us to identify markers for all and characterize morphology for many of them. Because ACs are known to display remarkable heterogeneity in neurotransmitter phenotype, we systematically analyzed expression of neurotransmitter biosynthetic enzymes and neuropeptide precursors, providing evidence for the presence of at least 20 small molecule or peptide transmitters in ACs, with potential use of multiple transmitters in the majority of them. We also analyzed transcriptional relationships among types and identified transcription factors expressed by closely related types. They include *Meis2* and *Tcf4*, expressed by most GABAergic and glycinergic types, respectively.

Combined with results from other classes, our inventory of ACs provides what we believe to be a nearly complete mouse retinal cell atlas, comprising approximately 140 cell types. Thus, at least in this respect, the retina is about as complicated as any other part of the brain.

## MATERIALS AND METHODS

### Animals

Animals were used in accordance with NIH guidelines and protocols approved by the Institutional Animal Care and Use Committee (IACUC) at Harvard University. The following knock-in and transgenic mouse lines were obtained from Jackson Laboratories: *Chx10-cre-GFP* (Rowan and Cepko, 2004; Stock No:005105; *Chx10* is now named *Vsx2*), *Slc17a7-IRES2-Cre* (Harris et al., 2014 Stock No: 023527), *Cck-IRES-Cre* (Taniguchi et. al, 2011; Stock No: 012706), *Penk-IRES2-Cre* (generated at the Allen Institute; JAX Stock No: 025112), *Sst-IRES-Cre* (Stock No: 013044) and *Thy1-mitoCFP-P* (Misgeld et al, 2007; Stock No: 007967). *Contactin 5-lacZ* and *Contactin 6-lacZ* lines were from Sudo and colleagues (Li et al., 2003; Takeda et al., 2003) via Julia Kaltschmidt and Thomas Bourgeron, respectively. *NeuroD6-cre* knock-in mice (Goebbels et al., 2006) were obtained from K. Nave via L. Reichardt. The *Gbx2-CreERT2-IRES-GFP* line (Chen et al., 2009) was a generous gift from James Y. H. Li (University of Connecticut) via Chinfei Chen’s lab (Harvard Medical School). The *Ptf1a-cre* (Kawaguchi et. al, 2002) line was obtained from Lisa Goodrich (Harvard Medical School), The *Thy1-STP-YFP* Line 15 was generated in our laboratory (Buffeli et. al, 2003; JAX Stock No:005630). Some tissue was obtained from mice analyzed in a previous study (Martersteck et al., 2017). *Ptf1a-cre* was maintained on a CD1 background (Charles River Stock No. 022). All other mutants were maintained on a C57BL/6J background (JAX Stock No. 000664).

### Cell Sorting and Single-Cell Sequencing

Post-natal day 19 (P19) *Chx10-cre-GFP* animals were euthanized by intraperitoneal injection of Euthasol. Eyes were removed and retinas dissected in oxygenated Ames solution. Retinas were incubated at 37°C for 10 minutes in a papain solution followed by trituration in an ovomucoid solution to quench papain activity and generate a single cell suspension. Cells were centrifuged at 450 x g for 8 minutes and the pellet was resuspended in Ames+4% BSA with rat anti-mouse CD133-APC (eBioscience) and rat anti-mouse CD73-PE (eBioscience). Following incubation for 15 minutes at room temperature, cells were washed with Ames + BSA, centrifuged again, and resuspended at a concentration appropriate for flow cytometry. Samples were sorted on a MoFlo Astrios cell sorter (Beckman) and cells triple negative for GFP, APC, and PE but positive for the cell viability marker Calcein Blue were collected. They were then processed according to the 10X Genomics v2 Chromium Single Cell 3’ Reagent Kit (10X Genomics; Zheng et al., 2017). Briefly, single cells are partitioned into oil droplets containing single oligonucleotide-derivatized beads followed by cell lysis, barcoded reverse transcription of RNA, amplification, shearing, and attachment of 5’ adaptor and sample index oligos. Libraries were sequenced on the Illumina HiSeq 2500 (Paired end reads: Read 1, 26bp, Read 2, 98bp).

Adult amacrine cells were collected and sequenced as part of unrelated projects (Tran et al., 2019; I.B., I.E.W., N. M.T., and J.R.S., unpublished).

### Computational Methods

We analyzed scRNA-seq data following the pipeline detailed in Peng et al. (2019). Briefly, sample demultiplexing was performed with cellranger mkfastq function (version 2.1.0; 10X Chromium), and reads were aligned to the reference genome mm10 v1.2.0 from cellranger refdata using the cellranger count function with the option --force-cells=6000. Clustering was performed to stratify cells into major classes using defining class-selective markers, and then to cluster amacrine cells into putative types. Subsequent steps were as follows: 1) A threshold of 600 genes detected per cell was applied to filter out low quality cells and debris. 2) Gene expression matrix was calculated as UMI count matrix first normalized by total number of UMI for each cell and multiplied by the median UMI count per group, and lastly log transformed after adding 1 (Shekhar et al., 2016). 3) Highly variable genes were identified by the method of Pandey et al, (2018). 4) Batch correction was performed on the highly variable genes expression matrix using a linear regression model in the Seurat package (https://satijalab.org/seurat/). 5) Principal component (PC) analysis was applied and significant PCs estimated based on Tracy–Widom theory (Patterson et al., 2006) were used for further analysis of clustering. 6) Data were partitioned into clusters of transcriptionally related cells using the Louvain algorithm with the Jaccard correction (Shekhar et al., 2016). 7) A dendrogram was built on the expression matrix of HVGs for the assigned clusters to reveal their overall transcriptomic similarity. 8) Clusters closest to each other on the dendrogram were assessed for differential expression (DE), and iteratively merged if no more than six DE genes were found (log fold change >1 and adjusted p value <0.001). DE tests were performed using the R package ‘MAST’ (Finak et al., 2015). 9) Because some contaminants (i,e,, non-ACs or doublets) became evident only following clustering, we reexamined the clustered data to remove them. Criteria for doublets included having an increased number of transcripts per cell and having combined expression of canonical markers from different cell classes. 10) We retested for doublets among amacrine cells using the R package ‘DoubletFinder’ (McGinnis, et al. 2019) with the default parameter of 7.5% as expected doublet rate. We found that ∼55% of cells in clusters 16, and 60 could be doublets, although we have no way of verifying this possibility. As a second test for amacrine-amacrine doublets, we asked whether each cluster expressed genes at far higher levels than any other cluster. Although there are some variations among clusters, C16 was suspect, but all others appeared to be authentic. 11) To compare types between datasets, we trained multi-class classifiers (Xgboost algorithm; Tianqi Chen, 2016) using the R package ‘xgboost’ and assigning matches as detailed in Peng et al. (2019). 12) For visualization purposes only, dimensionality was further reduced to 2D using Uniform Manifold Approximation and Projection (UMAP).

### Histology

Following euthanasia with Euthasol mouse eyes were fixed in 4% paraformaldehyde in 1X phosphate buffered saline (PFA/PBS). In most cases, PFA/PBS was administered by transcardiac perfusion followed by eye removal and a 20 min post-fix in PFA/PBS. Alternatively, eyes were removed immediately and fixed in PFA/PBS for 90 min on ice. Eyes were then rinsed with PBS and the retina dissected out. Retinas to be sectioned were incubated in 30% sucrose in PBS overnight at 4°C, after which they were embedded in tissue freezing medium, frozen in dry ice, and stored at -80°C until processing. Retinas were cryosectioned at 20-25μm and air dried. The sections were rehydrated in PBS, incubated in 5% normal donkey serum, 0.3% Triton X-100 in PBS for 1 hour, incubated with primary antibodies overnight at 4°C, washed in PBS, incubated with secondary antibodies for 2 hours at room temperature, washed again in PBS, allowed to dry, mounted with Vectashield (Vector Lab), and cover slipped.

For whole mounts, retinas were blocked in 5% donkey serum, 0.3% Triton X-100 in PBS for 3-14 hours and incubated in primary antibody for 5-7 days at 4°C. Retinas were then washed in PBS and incubated overnight in secondary antibody. Finally, retinas were washed in PBS, flat-mounted on cellulose membrane filters (Millipore), cover slipped with Fluoro-Gel (Electron Microscopy Sciences), and sealed with nail polish.

The following antibodies were used rabbit and chicken anti-GFP (1:2000, Millipore; 1:1000, Abcam); goat anti-choline acetyltransferase (ChAT) (1:500, Millipore); goat anti-vesicular acetylcholine transporter (VAChT) (1:500, Santa Cruz Biotechnology); mouse anti-Tfap2b (1:200, DSHB); rabbit anti-Rbpms (1:500, Abcam); guinea pig anti-Rbpms (1:1000, Phosphosolutions); goat anti-Vsx2 (1:300, Santa Cruz); rabbit anti-Vglut3 (1:1000, Synaptic Systems); rabbit anti-Ebf3 (1:2000, Millipore); rat anti-CD140a-PE (1:100, Thermo Fisher); rabbit anti-Nfix (1:1000, Thermo Fisher); rat anti-Somatostatin (1:500, Millipore); rabbit anti-βGAL (1:5000, in house); rabbit anti-Ppp1r17 (1:1000, Atlas Antibodies); rabbit anti-Neuropeptide Y (1:1000; Abcam); rabbit anti-Ghrh (1:500; Abcam), goat anti-Glyt1 (1:10000, Chemicon), mouse anti-GAD 65/67 (1:500, DSHB), rabbit anti-GAD65/67 (1:1000, Millipore, AB1511), mouse anti-Pax6 (1:500, DSHB), mouse anti-Meis2 1A1 (1:100, DSHB); guinea pig anti-Prdm8 (1:2000, kind gift from Sarah E Ross lab, University of Pittsburgh), guinea pig anti-Lhx9 #1342 (1:5000, kind gift from Jane Dodd lab, Columbia University Medical Center), mouse anti-Tfap2c (1:500, Sigma-Aldrich), rabbit anti-Calbindin (1:2000, Swant), mouse anti-Calretinin (1:5000, Millipore), rabbit anti-Neurod2 (1:500, Abcam), rabbit anti-Tbr2 (1:500, Abcam), rabbit anti-TCF4 (1:200, Proteintech). Secondary antibodies were conjugated to Alexa Fluor 488 (Invitrogen), Alexa Fluor 568 (Invitrogen), or Alexa Fluor 647 (Jackson ImmunoResearch) and used at 1:1000. Nuclei were stained with ToPro Cy5 (1:5000, Thermo Fisher).

To sparsely label ACs, we injected mice from Cre-expressing lines listed above with a cre-dependent virus, *AAV9-EF1a-BbTagBY* (Cai et al., 2013; Addgene #45185-AAV9), *AAV9-EF1a-BbChT* (Cai et al., 2013; Addgene #45186-AAV9) or *AAV9-CAG-tdTomato* (Addgene #51503-AAV9). Intravitreal injections were performed as described in Tran et al. (2019). Animals were euthanized 3-4 weeks after injection and retinas processed as above.

### Probe generation for *in situ* hybridization

To generate probes, RNA was extracted from P19 C57BL/6J mouse retina and reverse transcribed as described in Laboulaye et al. (2018). Probes were generated from cDNA by PCR using Q5 Polymerase and the following sets of primers.

*Car3* (901bp)

Forward: 5’-ATCTTCACTGGGGCTCCTCT-3’

Reverse: 5’-gaaattaatacgactcactatagggCGCATACTCCTCCATACCCG-3’

*Kit* (1017bp):

Forward: 5’-TGGTCAAAGGAAATGCACGA-3’

Reverse: 5’-gaaattaatacgactcactatagggTCTTCTTAGCGTGACCAG-3’

*Slc17a7* (1028bp):

Forward: 5’-CGGATACTCGCACTCCAAGG-3’

Reverse: 5’-gaaattaatacgactcactatagggTTCCCTCAGAAACGCTGGTG-3’

PCR products were separated by gel electrophoresis, after which bands of the expected size were column purified and sequenced. Confirmed PCR templates were then transcribed by T7 RNA polymerase (Roche) with a DIG-UTP nucleotide mix (Roche). Resulting products were precipitated overnight at -80°C in mixture of TE buffer, LiCl, and EtOH. The following day, samples were centrifuged. Pellets were then washed with 70% EtOH and resuspended with 1:1 formamide/water. Finally, probes were run on agarose gel to ensure expected size.

### Fluorescent *in situ* hybridization (FISH)

Tissue was collected and prepared with RNase-free reagents and sectioned as described above. Slides were processed as described in Tran et al. (2019). Briefly, slides were fixed in 4% PFA for 10 minutes and then rinsed 2 x 5 min PBS with 0.1% Tween-20 (PBT). Slides were then digested in a Proteinase-K solution of 0.5 µg/ml for 3 minutes, washed 2 x 5 min in PBT, and fixed again in 4% PFA for 5 min. Slides were then washed 2 x 5 min in PBT, incubated in acetylation solution (0.1M Triethanolamine + 0.25% acetic anhydride) for 5 min. Slides were then washed 2 x 5 min in PBT and incubated in pre-hybridization solution for 1 hour at room temperature. Digoxigenin (DIG) labeled probes were denatured for 5 minutes at 85°C and added to slides at 1:200 dilution. Slides were then cover slipped and incubated overnight at 65°C.

On the second day, slides were washed 2x in pre-hybridization solution, 2x in 2X saline sodium citrate (SSC): Prehybridization Solution, 2x in 2X SSC, and 2x in 0.2X SSC. All of these washes were conducted for 30 minutes each at 65°C. Slides were then washed 2x in Maleic Acid Buffer containing 0.1% Tween-20 (MABT) at room temperature and blocked in Heat Inactivated Sheep Serum (HISS)/MABT/blocking solution for 1 hour. Slides were then incubated overnight with anti-DIG-HRP antibodies (1:750).

On the third day, slides were washed 6 x 5 min in MABT, followed by 2 x 5 min in PBT. Signals were amplified with Cy3-tyramide (1:200) for 1 hour (TSA-Plus System; Perkin-Elmer Life Sciences, MA). Slides were rinsed 6×5 minutes in PBT and 2×5 minutes in 1X PBS and then incubated in 3% donkey serum, 0.3% Triton-X in 1X PBS for 30 minutes, followed by primary antibody incubation overnight at 4°C.

On the last day, slides were washed 3 x 5 min in 1X PBS. Slides were then incubated in secondary antibodies for 2 hours at room temperature, washed 3x 5 min in 1X PBS, dried, cover-slipped, and sealed.

Images were acquired on an Olympus-FV1000 Confocal Microscope. We used ImageJ (NIH) software to analyze confocal stacks and generate maximum intensity projections.

### Experimental Design and Statistical Analysis

Statistical methods for analysis of RNA-seq data are detailed above (see “Computational methods”). To quantify immunostaining combinations shown in Figure 9, confocal images were analyzed in ImageJ. ≥4 images from ≥2 retinas were analyzed for each marker combination. Custom ImageJ macros were used to place circular regions-of-interest (ROIs) (3.44 µm in diameter) over all cell somas and nuclei that were positive for at least one marker. Fluorescent intensity was measured for each marker in each ROI in single Z-slice images. Fluorescent values for ROIs from each marker were minimum subtracted and normalized to the maximum value for plotting.

### Code and data accessibility

Submission of all the raw and processed datasets reported in this study has been initiated to the Gene Expression Omnibus (GEO) with accession number GSE xxx (private until publication). The single cell data can be visualized in the Broad Institute’s Single Cell Portal at https://singlecell.broadinstitute.org/single_cell/study/xxxxxxx (private until publication).

## RESULTS

### Transcriptomic separation of amacrine cells into 63 clusters

In previous studies aimed at generating BC and RGC atlases, we enriched cells using class-specific transgenic markers, Vsx2/Chx10 and VGlut2, respectively (Shekhar et al., 2016; Tran et al., 2019). We were unable to find a suitable class-specific marker for ACs, and therefore adopted a strategy of selective depletion. We dissociated retinas from postnatal day (P)18-19 *Vsx2-GFP* mice, in which BCs and Müller glia are labeled (Rowan and Cepko, 2004; Shekhar et al., 2016), and labeled rod and cone photoreceptors with fluorophore-conjugated antibodies to CD73 and CD133, respectively (Lakowski, 2011; Peng et al., 2019). We isolated GFP/CD73/CD133 triple negative cells by FACS and used droplet-based scRNA-seq (Zheng et al., 2017) to obtained high quality transcriptomes from 55,287 cells. We then divided them into classes by expression of canonical markers (Peng et al., 2019; Figure 1C). ACs were defined as cells that were positive for the transcription factor *Pax6* and the vesicular inhibitory amino acid transporter *Slc32a1*, and negative for markers of other classes; they comprised 58.8% of the population (Figure 1D) or a total of 32,523 ACs. Unsupervised analysis divided the ACs into 63 clusters, each being a putative cell type (Figure 1E). They ranged in abundance from 0.1-7.7 percent of all ACs (Figure 1F). Since ACs comprise 7-10% of retinal cells (Jeon et al., 1998; Macosko et al., 2015), individual types comprise ∼0.01-1% of all retinal cells.

Before proceeding further, we performed three tests to assess the possibility that our collection protocol had excluded AC types. First, we immunostained “uncollected” cells from the FACS isolation procedure with the AC marker TFAP2B, and found negligible numbers of positive cells, indicating that few if any AC types were GFP-, CD73- or CD133-positive (data not shown). Second, we trained a classifier using a supervised machine learning algorithm, XGBoost (Tianqi Chen, 2016) to match the 21 groups identified from a collection that included no enrichment or depletion steps (Macosko et al., 2015) to the 63 clusters in the current dataset (Figure 2A). As expected, in many cases, a single cluster in the smaller dataset (∼4.4k ACs) mapped to multiple clusters in our larger dataset, indicating improved resolution in distinguishing cell types. Importantly, however, no clusters from Macosko dataset were left unmatched in our data. Finally, we asked whether some types might emerge later than P19. To this end, we queried a set of 5,347 ACs collected from P56 mice in the course of unrelated studies (Tran et al., 2019; I.B, I.E.W., N.M.T. and J.R.S., unpublished). As with the Macosko AC data, the P56 AC data was underpowered to resolve every type but was sufficient to resolve 20 clusters. Again, all P56 AC clusters mapped to one or more P19 clusters, and in no case was a P56 cluster unmatched in the P19 dataset, as might occur if further diversification occurred following P19 (Figure 2B). Together, these results support the idea that our dataset includes most if not all AC types that comprise >0.01 percent of retinal cells.

**Figure 2.**
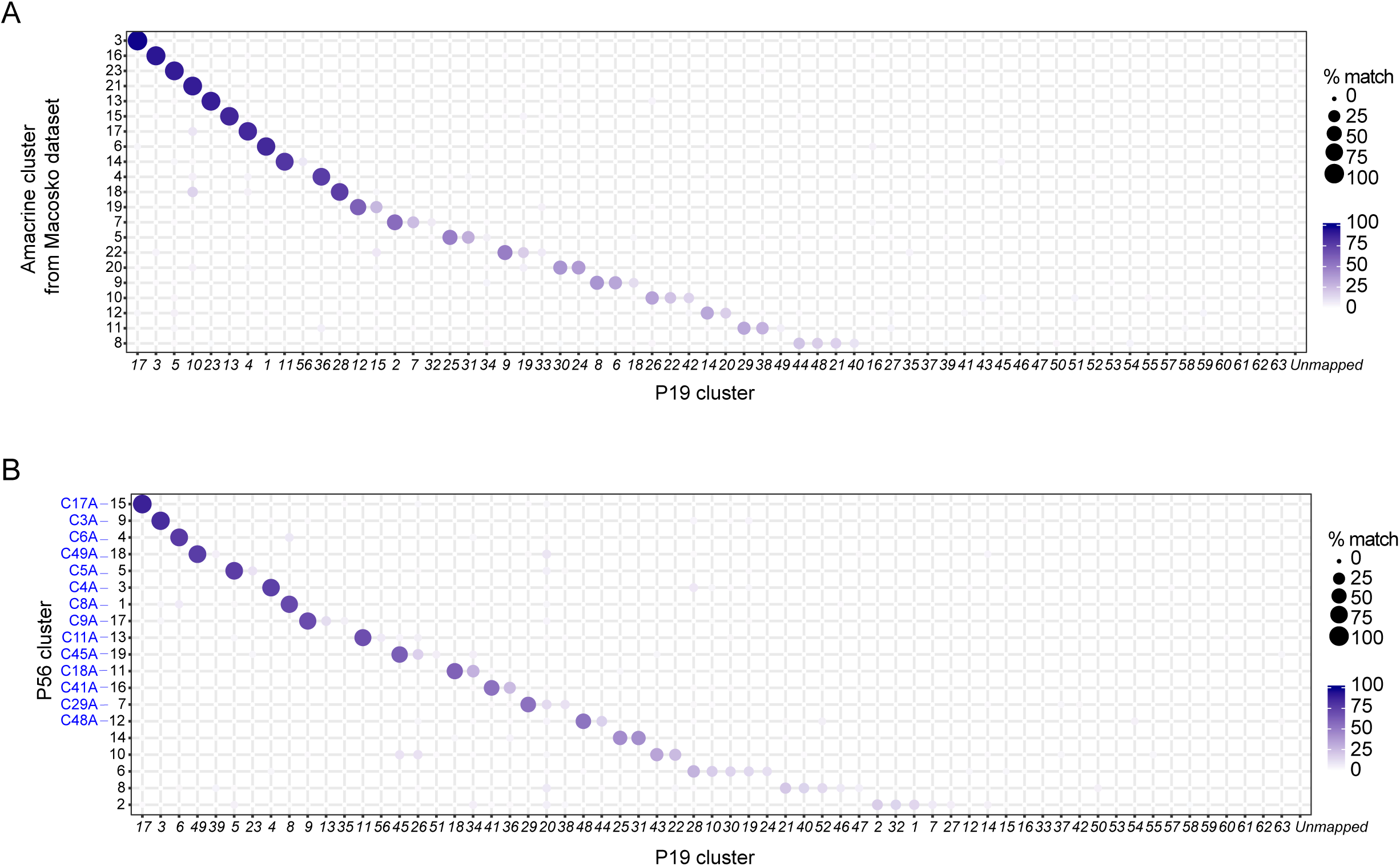
No additional AC types detected in two other datasets. A. Comparison of AC clusters obtained with selective depletion of other cell classes (this study) or without (Macosko et al., 2015). Transcriptional correspondence is depicted as a matrix in which circle diameter and color indicate the percentage of cells in a given AC type from the new data (x axis) that are assigned to a particular type from Macosko et al (y axis) by a classification algorithm (xgBoost) trained on the P19 data. For the y axis, clusters were ordered by degree of match to a single P19 cluster. While some AC types were not represented in the Macosko et al. data, all types detected by Macosko et al., were represented in the new dataset. B. Comparison of AC clusters obtained in this study with those sampled from P56 retina. While some AC types were not represented in the P56 data, all P56 types were represented in the new dataset. We re-named P56 clusters in which more than 50% cells mapped to single P19 clusters with their P19 counterparts’ cluster ID plus “A”.

### Correspondence between clusters and AC types

To match molecularly defined clusters to authentic AC types, we began by identifying differentially expressed (DE) genes for each cluster. Most clusters could be uniquely identified by expression of a single DE gene (Figure 3A) and the others by a combination of two DE genes. These and other DE genes allowed assignment of several clusters to previously identified AC types (Table 1). They included starburst ACs (C17: *Chat*; Vaney et al., 2012), Aii ACs (C3: *Gjd2, Prox1, Dab1, Nfia, Dner*; Hansen et al., 2005; Rice and Curran, 2000; Perez de Sevilla Müller et al., 2017; Keeley and Reese, 2018), SEGs (C4: *Satb2, Ebf3*, and Glyt1 [*Slc6a9*]; Kay et. al, 2011), VG3 ACs (C13: VGlut3 [*Slc17a8*]; Haverkamp and Wässle, 2004; Johnson et al., 2004; Krishnaswamy et al., 2015); and A17 ACs (C6: *Prkca* (PKCα), *Sdk1, Calb2* negative, Dab1-negative; Grimes et al., 2010; Puthussery and Fletcher 2007; Yamagata and Sanes, 2019). Other types expressed neuropeptides known to mark specific AC types; they include *Cck* (C10,C17,C18,C34; Firth et al., 2002), *Vip*, (C22,C26,C47; Park et al., 2015; Akrouh and Kerschensteiner, 2015; Pérez de Sevilla Müller et al., 2019), *Crh* (C37; Park et al., 2018) and *Penk* (C35,C59,C63; Chen et al., 2013). For some of these types, we were able to validate new markers, such as *Car3* for VG3 cells (Figure 3B,C).

**Table 1.** Summary of AC types.

| Cluster | GABAergic/Glycinergic | # cells profiled | Type or key marker |
| --- | --- | --- | --- |
| 1 | GABAergic | 2505 |  |
| 2 | GABAergic | 2190 |  |
| 3 | Glycinergic | 1897 | Aii |
| 4 | Glycinergic | 1618 | SEG |
| 5 | GABAergic | 1396 |  |
| 6 | GABAergic | 1353 | A17 |
| 7 | GABAergic | 1216 |  |
| 8 | GABAergic | 1029 |  |
| 9 | Glycinergic | 800 |  |
| 10 | Neither | 754 | nGnG-2; CCK |
| 11 | GABAergic | 726 |  |
| 12 | Glycinergic | 721 |  |
| 13 | Glycinergic | 707 | VG3 |
| 14 | GABAergic | 698 |  |
| 15 | Glycinergic | 663 |  |
| 16 | Both | 638 |  |
| 17 | GABAergic | 637 | SAC |
| 18 | GABAergic | 621 | CCK |
| 19 | Glycinergic | 588 |  |
| 20 | GABAergic | 585 |  |
| 21 | GABAergic | 574 |  |
| 22 | GABAergic | 559 | VIP |
| 23 | GABAergic | 533 |  |
| 24 | Neither | 480 | nGnG-1 |
| 25 | GABAergic | 480 | CA-II |
| 26 | GABAergic | 470 | VIP |
| 27 | GABAergic | 468 |  |
| 28 | Glycinergic | 462 |  |
| 29 | GABAergic | 459 |  |
| 30 | Neither | 435 | nGnG-3 |
| 31 | GABAergic | 433 |  |
| 32 | GABAergic | 421 |  |
| 33 | Glycinergic | 360 |  |
| 34 | GABAergic | 348 | CCK |
| 35 | Glycinergic | 300 | PENK |
| 36 | Neither | 289 | nGnG-4 |
| 37 | GABAergic | 280 | CRH |
| 38 | GABAergic | 277 |  |
| 39 | GABAergic | 262 |  |
| 40 | GABAergic | 257 |  |
| 41 | GABAergic | 230 |  |
| 42 | GABAergic | 221 |  |
| 43 | GABAergic | 220 |  |
| 44 | GABAergic | 210 |  |
| 45 | GABAergic | 206 | CA-I |
| 46 | GABAergic | 203 |  |
| 47 | GABAergic | 197 | VIP |
| 48 | GABAergic | 184 | nNOS |
| 49 | GABAergic | 175 |  |
| 50 | GABAergic | 159 |  |
| 51 | GABAergic | 130 | GHRH |
| 52 | GABAergic | 118 | nNOS |
| 53 | Both | 109 |  |
| 54 | GABAergic | 91 | nNOS |
| 55 | GABAergic | 89 |  |
| 56 | GABAergic | 86 | VG1 |
| 57 | GABAergic | 81 |  |
| 58 | GABAergic | 81 |  |
| 59 | GABAergic | 77 | PENK |
| 60 | GABAergic | 64 |  |
| 61 | GABAergic | 41 |  |
| 62 | Both | 34 |  |
| 63 | GABAergic | 28 | PENK, SST |

**Figure 3.**
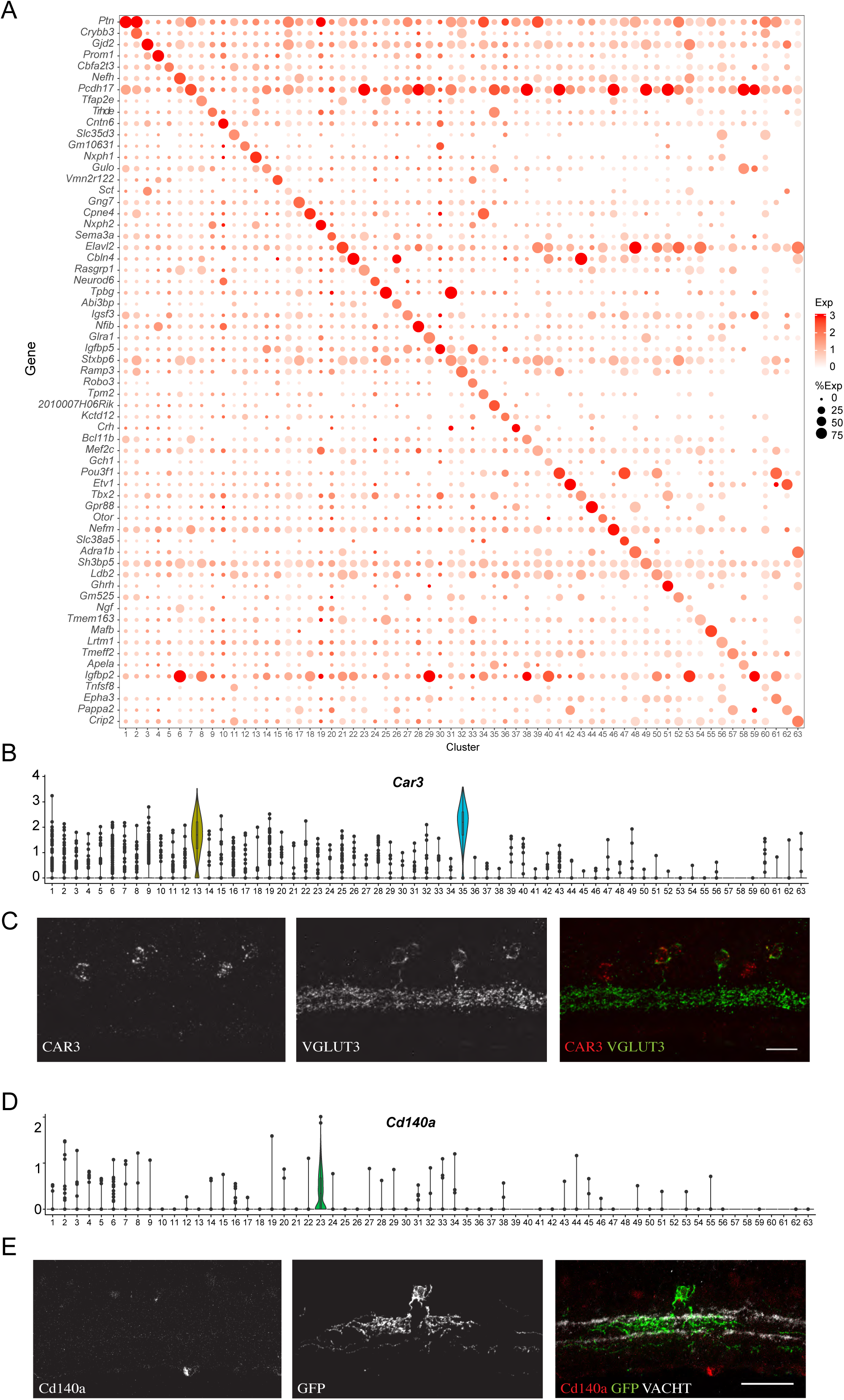
Molecular markers of AC types. A. Dot plots showing genes (rows) that uniquely mark AC clusters (columns). In this and subsequent figures, the size of each circle is proportional to the percentage of cells expressing the gene, and the color depicts the average transcript count in expressing cells. B. Violin plots representing expression of *Car3* in each of the AC types. Expression is highest in C35 (VG3-ACs) but also substantial in C13. Numbers above each type show percent of cells that expressed the gene. C. Retinal section doubly labeled for *Car3* (in situ hybridization) and VGlut3 (immunohistochemistry). VG3 ACs are *Car3*-positive. Scale bar is 20 µm. D. Violin plot showing selective expression of *Cd140a* in C23. E. Retinal section sparely labeled with GFP (Ptf1-cre x cre-dependent AAV) and CD140A antibody reveals morphology of C23 ACs. Scale bar is 40µm.

To characterize AC types that had not, to our knowledge, been studied previously, we combined fluorescent *in situ* hybridization or immunohistochemistry with sparse labeling to reveal cellular morphology. We used lines that express cre recombinase in subsets of ACs, and infected retinas with adeno-associated viral (AAV) vectors that express a fluorescent protein in a Cre-dependent manner. For example, immunostaining sections from an AAV-infected *Ptf1a-cre* line, which expresses broadly in newly born ACs (Fujitani et. al, 2006), showed that C23 (*Cd140a*-positive) is a medium-field AC type that stratifies in the center of the inner plexiform layer (IPL; Figure 3D,E). Additional examples are presented below. Thus, although our mapping of molecular to morphological types is not exhaustive we conclude that most or all of the AC clusters identified in our dataset correspond to authentic cell types.

As a further test of these assignments, we examined clusters from the adult (P56) dataset that matched with a single one of the 63 AC clusters. For convenience, we renumbered these clusters so they correspond to their P19 counterpart (Figure 2B), referring to them as C3A for P56 cluster 9, C4A for P56 cluster 3 and so on. They included starburst, A17, SEG, and Aii ACs as well as *Nos1*- and *Cck* positive clusters. In each case, most or all of the defining markers identified at P19 were retained at P56 (Figure 4). A notable exception was a catecholaminergic type (C45, C45A), which is further examined below.

**Figure 4.**
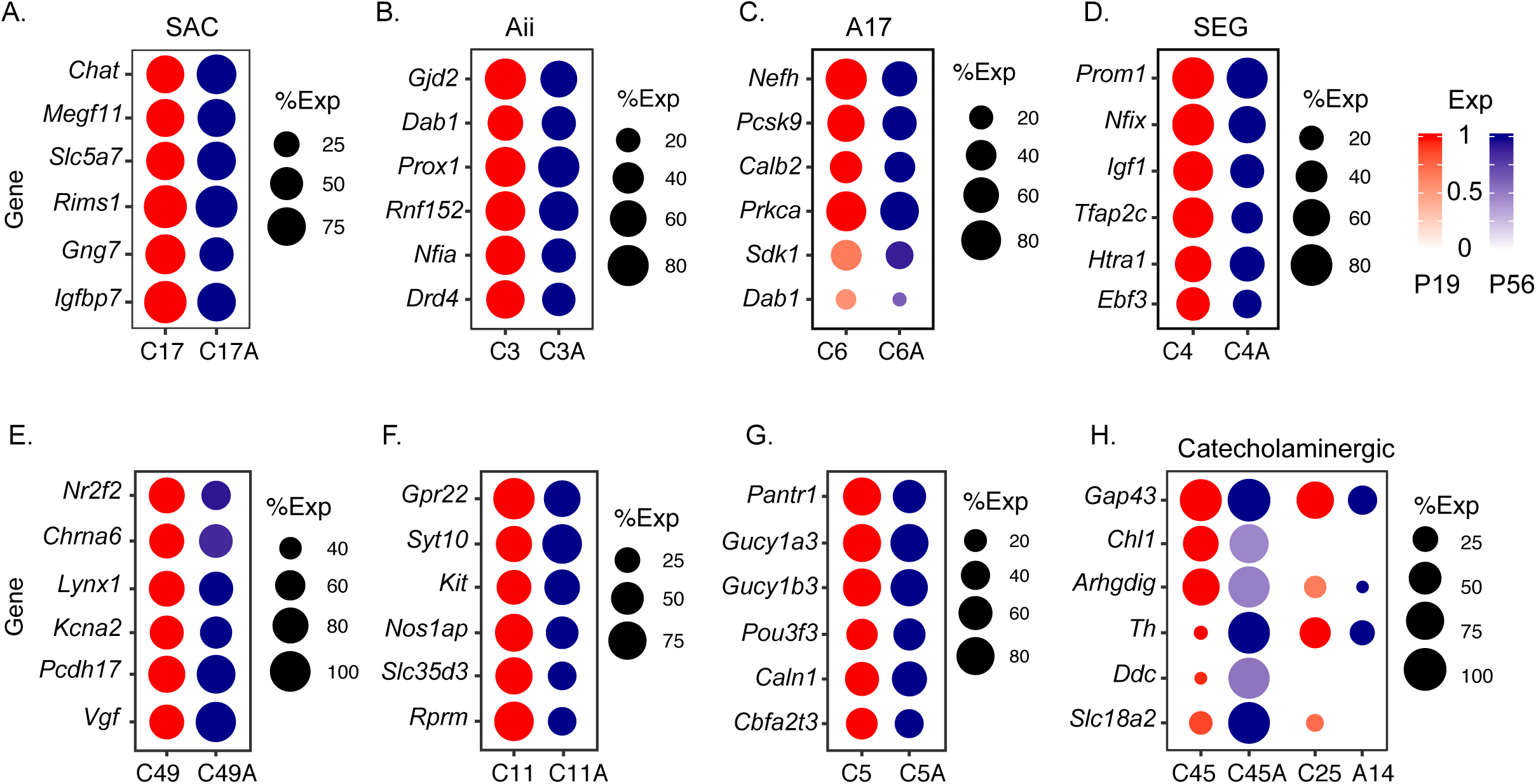
Expression of cell type-specific genes in selected types of ACs at P19 and in adults. Dot plots show marker genes for Starburst amacrine cells (C17; A), Aii ACs (C3; B), A17 ACs (C6; C), SEG ACs (C4, D), C49 (E), C11 (F), C5 (G) and CAI and CAII catecholaminergic ACs (C45 and C25;H), Most markers were maintained for most types. For catecholaminergic ACs, however, levels of dopamine synthetic enzymes (*Th, Ddc*) and the monoamine transporter (*Slc18a2*) were several-fold higher in adult than in P19 C45 ACs, suggesting that C45 is the CAI AC type. Adult cluster A14 is the closest match to the P19 C25 (Figure 2B).

### Neurotransmitters and neuromodulators

Most ACs are inhibitory neurons that use the neurotransmitter γ-aminobutyric acid (GABA) or Glycine. However, it has long been known that subsets of ACs also contain a variety of other small molecule and peptide neurotransmitters and neuromodulators (Karten and Brecha, 1983). Our AC atlas provided an opportunity to systematically characterize these subsets. For this analysis, we arranged the types by transcriptomic similarity based on hierarchical clustering, rather than frequency (Figure 5A), so that we could assess relationships among types that shared transmitters.

**Figure 5.**
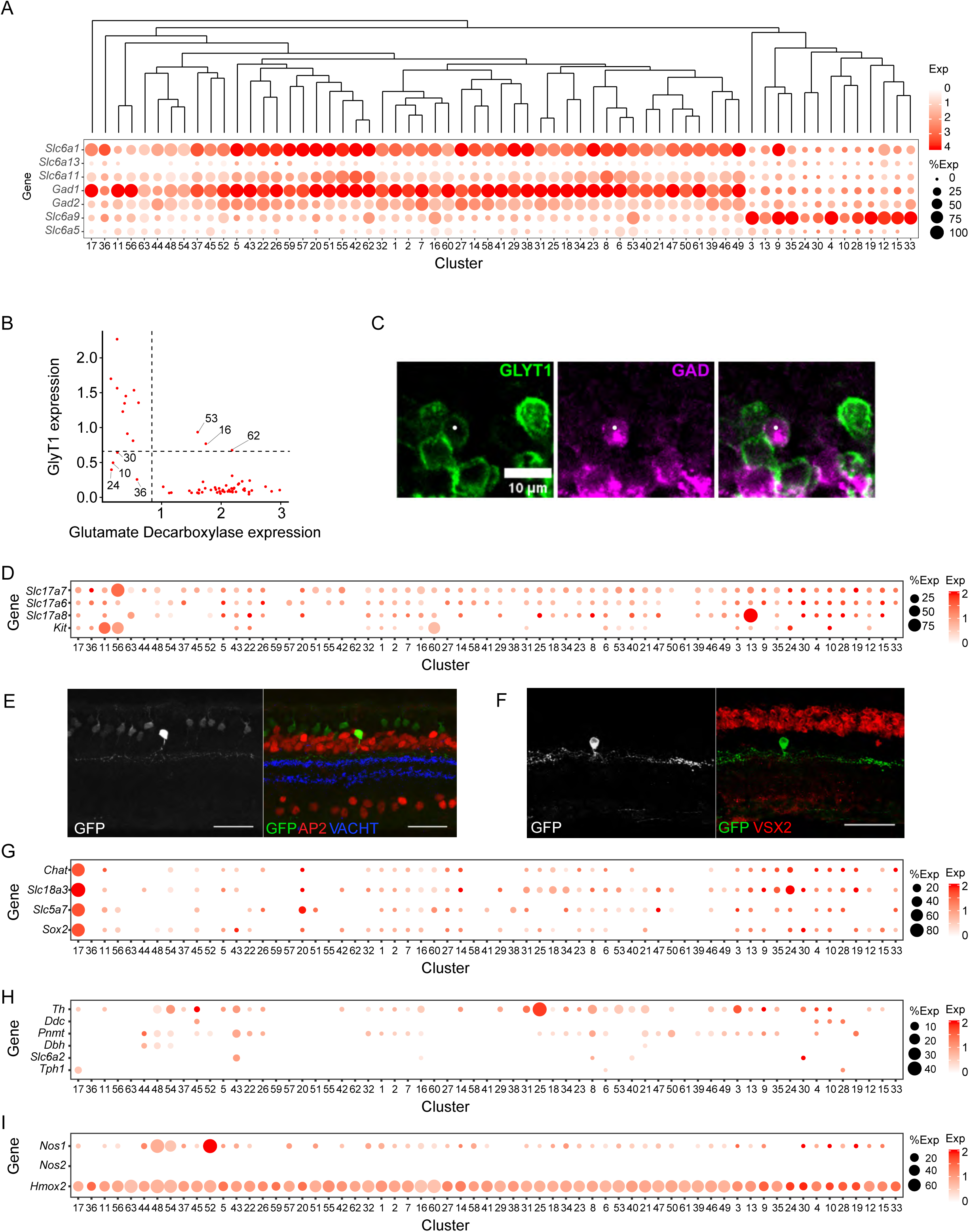
Evidence for multiple small molecule transmitters in ACs. A. Expression of GABAergic and glycinergic markers in ACs. *Slc6a1, Slc6a13* and *Slc6a11* encode GABA transporters type 1, 2 and 3, respectively. *Gad1* and *Gad2* encode two isoforms of glutamate decarboxylase. *Slc6a9* and *Slc6a5* encode vesicular glycine transporters 1 and 2, respectively. B. Expression of the most informative GABAergic (*Gad1* and *Gad2*) and glycinergic (*Slc6a9*) markers divides ACs into 4 groups: GABAergic (43 types), glycinergic (11 types), neither (nGnGs, 4 types) and potentially both GABAergic and glycinergic (3 types). C. Immunostaining of P22 retinas revealed sparsely populated ACs (white dot) that co-express GABAergic (GAD65/67) and glycinergic (GLYT1) markers. Scale bar, 10µm. D. Distinct AC clusters express glutamate transporters *Slc17a7* (VGlut1; VG1 ACs) and *Slc17a8* (VGlut3; VG3 ACs); none express *Slc17a6* (VGlut2). E-F. Characterization of VG1 ACs, labeled in *VGlut1-Cre* x *Thy1-STP-YFP* line 15 mice. VG1 ACs are AP2-positive (D) and Vsx2-negative (E). Scale bars are 40µm. G. A single cluster (C17) expresses the cholinergic markers choline acetyltransferase (ChAT), *Slc18a3* (Vesicular acetylcholine transporter, VACht) and *Slc5a7* (choline transporter). H. Expression of genes encoding synthetic enzymes for monoamine neurotransmitters. I. Two AC clusters express *Nos1* (nitric oxide synthetase type 1) and therefore could use NO as a transmitter.

#### GABA and Glycine

We examined expression of the GABA synthetic enzyme glutamate decarboxylase (*Gad1* and *Gad2*), and GABA transporters *Gat 1-3* (*Slc6a1, Slc6a11* and *Slc6a13*) as markers of GABAergic cells, and expression of glycine transporters *GlyT1* and *GlyT2* (*Slc6a9* and *Slc6a5*) as markers of glycinergic cells (Figure 5A). Most informative were the canonical markers *Gad1, Gad2* and *Glyt1*. As expected, expression of *Gad* (1+2) and *GlyT1* was mutually exclusive in most types, with 43 of the 63 types being GABAergic and 13 being glycinergic (Figures 5A,B). The GABAergic and glycinergic types were entirely restricted to different clades with the exception of starburst ACs (SACs; C17), which were distinct from either clade (see Discussion).

The remaining 7 types had unconventional neurotransmitter expression patterns (Figure 5B). Four types (C10, 24, 30 and 36) expressed markedly lower levels of both GABAergic and glycinergic markers, and likely represent “non-GABAergic, non-Glycinergic” (nGnG) ACs (Kay et al., 2011; Macosko et al., 2015); further analysis of these types is presented below.

The three remaining types (C16, 53, and 62) expressed high levels of both *Gad* (1+2) and *GlyT1*, raising the possibility that they use both transmitters. We asked whether clusters might be composed of “doublets,” arising from co-occupancy of a single microbead by two cells. This may indeed be the case for C16, as judged by a doublet-detecting algorithm but is unlikely for the others (see Methods). Moreover, we observed a sparse population of ACs that were positive for both GAD and GlyT1 by immunostaining (Figure 5C).

Three of the nGnG types were members of the glycinergic clade, while the fourth nGnG type and all of the putative GABA+glycine types were members of the GABAergic clade. GABAergic and glycinergic types include both abundant and rare types (Figure 1F), whereas the dual and nGnG types were all rare (<2% of ACs per type). Overall, the GABAergic, glycinergic, nGnG, and dual ACs comprise *∼67%, 25%, 6%*, and *2%*, respectively, of all ACs.

#### Glutamate

One well-studied AC type, VG3, expresses *VGlut3* as well as *Glyt1*, and is capable of both excitatory glutamatergic and inhibitory glycinergic transmission, likely at different synapses (Lee et al., 2016; Tien et al., 2016). We assessed expression of all three vesicular glutamate transporters. *VGlut1 (Slc17a7), VGlut2 (Slc17a6)*, and *VGlut3 (Slc17a8)* (Figure 5D). *VGlut3* expression was confined to the VG3 type and *VGlut2*, an RGC marker, was not detectably expressed. Surprisingly, we found one rare type, C56 (0.3% of all ACs), that expressed *VGlut1*, implying the existence of a second glutamatergic AC type. These cells, which we call VG1 ACs, expressed GABAergic but not glycinergic markers, and could potentially mediate dual excitatory glutamatergic and inhibitory GABAergic transmission.

We labeled these cells using a *VGlut1-Cre* line crossed to a cre-dependent reporter (*Thy1-STP-YFP*; Buffelli et al., 2013). In addition to bipolar cells, all of which are VGlut1-positive, and a rare VGlut1-positive RGC type (Tran et al., 2019), we detected VGlut1-positive cells that were ACs by the criteria that they were positive for the AC marker TFAP2B and negative for the bipolar marker VSX2 (Figure 5E,F). VG1 ACs laminated in S1 of the IPL (we divide the IPL into 5 sublaminae, with S1 abutting the inner nuclear layer and S5 abutting the ganglion cell layer) and have narrowly stratified but widely ramifying dendrites, which, consistent with their expression of *Gad*, is more characteristic of GABAergic (wide-field) than glycinergic (narrow-field) ACs.

#### Acetylcholine

Starburst ACs (C17, *Chat* positive) are intensively studied cholinergic ACs. We assessed expression of *Chat*, the vesicular acetylcholine transporter (*VAChT, Slc18a3*) and the high affinity choline transporter, *Slc5a7*. All were expressed selectively by C17 (Figure 5G), supporting the idea that starburst ACs are the only cholinergic ACs. ON and OFF starburst ACs are molecularly distinct in neonates (Peng et al., 2020) but differences between these closely related subtypes were no longer detectable by P19.

#### Monoamines

Monoamine neurotransmitters include dopamine, norepinephrine, epinephrine, serotonin, tyramine, tryptamine and histamine. We assessed expression of their synthetic enzymes (*Th, Ddc, Pnmt, Dbh, Tph1, Tph2*; Figure 5H) Tyrosine hydroxylase (*Th*), which generates DOPA from tyrosine, was expressed at high levels in C25, a GABAergic type and at lower levels in several other groups, consistent with evidence for at least two dopaminergic AC types, which vary in TH levels (Zhang et al., 2007; Vuong et al., 2015).

Surprisingly however, the other enzyme required for synthesis of dopamine, DOPA decarboxylase (Ddc; generates dopamine from DOPA) was not detectably expressed in C25, but traces of expression were detected in three clusters with low levels of TH (C4, 10 and 45). To understand this result, we examined expression of monoamine synthetic enzymes in adult ACs and found heterogeneity between clusters expressing TH (Figure 4H). C45A expressed highest levels of both TH and Ddc, marking it as the CAI AC. This cluster also selectively expressed connexin 45 (*Gja1*) which has been reported to label cells with a morphology characteristic of CAI ACs (Theofilas et al., 2017). C25A expressed lower levels of TH and no detectable Ddc, consistent with it being the CAII AC, in which dopamine and TH have been difficult to detect (Vuong et al., 2015). Additional molecular markers that may serve to differentiate between CAI and CAII ACs include Chl1, and Arhgdig. Other enzymes involve in synthesis of monoamine neurotransmitters were not expressed at significant levels by ACs at either P19 or P56 (Figure 5 and data not shown).

#### Gasotransmitters

Three gases have been implicated as neurotransmitters, NO, CO and H2S (Boehning and Snyder, 2003). Three AC types, C48, C52 and C54 expressed high or moderate levels of Nitric Oxide Synthase (*Nos1*; Figure 5I), consistent with previous reports of at least two Nos-positive AC types (Pang et al., 2010; Jacoby et al., 2018). Heme oxygenases (*Hmox1, Hmox2*) generate CO, with Hmox2 thought to be responsible for generating CO used as a neurotransmitter. *Hmox2* were broadly expressed by ACs (Figure 5H) whereas cystathionine γ-lyase (*Cth*) and cystathionine β-synthase (*Cbs*), which synthesize H2S, were not expressed at high levels by any ACs. Thus, we find no evidence for selective synthesis of CO or H2S by specific AC types.

#### Neuropeptides

We next mapped expression of genes encoding neuropeptides, which support a wide range of neuromodulatory and signaling roles (Figure 6A and Supplementary Figure 1A). In addition to those mentioned above, they included, Neuropeptide Y (*Npy*), Cocaine and amphetamine regulated transcript (*Cartpt*), Tachykinin (*Tac1*), Somatostatin (*Sst*), B-endorphin (*Pomc*), and Galanin (*Gal*). Neuropeptides were expressed by ACs from all small molecule neurotransmitter classes. We also observed expression of several neuromodulators previously studied in gut, brain, and other tissues but not, to our knowledge in retina, including Angiotensin, Calcitonin, Somatomedin and Ghrelin. In contrast, we observed no significant expression of 45 other peptides (Supplementary Figure 1).

**Figure 6.**
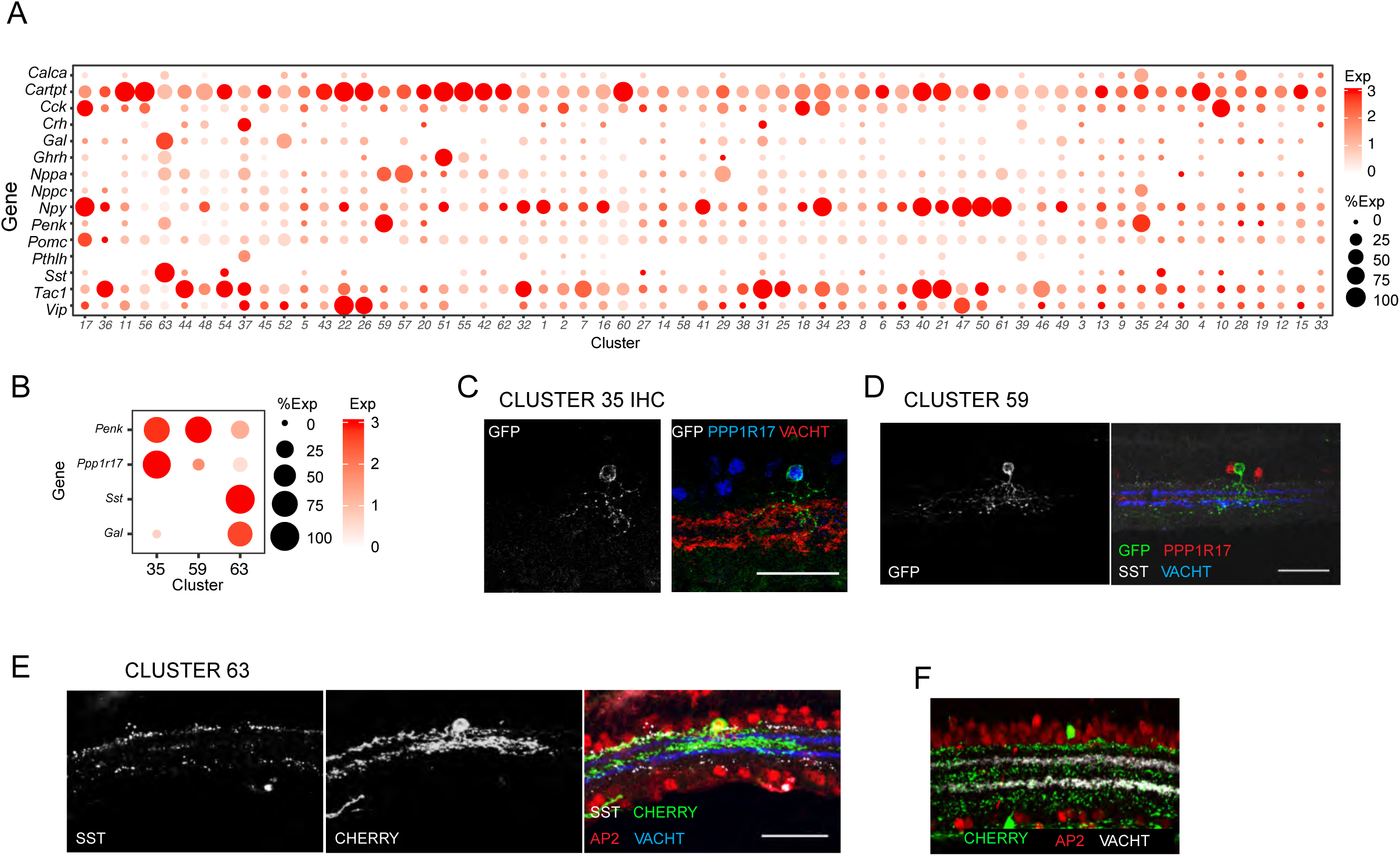
Many AC types express genes encoding neuromodulatory peptides. A. Dot plot showing peptides selectively expressed by one or a few AC clusters. (See Figure S2 for expression of other peptides.) B. Markers that distinguish three AC types that express the neuropeptide Penk. C. Penk-ACs were visualized by injection of a Cre-dependent AAV reporter into *Penk-cre* or *Sst-cre* mice. (C) C35 ACs were identified by co-expression of Penk and Ppp1r17. D. C59 ACs were both Sst- and Ppp1r17-negative. (E,F). C63 ACs were identified by co-expression of Penk and Sst (D) or co-expression of Sst and Tfap2b. Scale bar is 20µm in (C) and 40µm in (D-F).

Several neuropeptides were expressed by multiple AC types. For one of them, proenkephalin (encoded by the *Penk* gene), we combined immunohistochemistry with sparse labeling to assess morphology of individual types. *Penk* is expressed at highest levels by C35, C59 and C63 (Figure 6B). These types were distinguished by selective expression of *Ppp1r17* and *Car3* (C35), or *Sst* and *Gal* (C63); C59 express none of these genes. We marked and characterized *Penk*-positive cells by injecting a Cre-dependent AAV reporter into a *Penk-Cre* mouse line. C35 ACs (Ppp1r17+) are narrow-field ACs; C59 ACs (Sst- and Ppp1r17-) are medium-field; and C63 ACs, (Sst+) are widefield ACs with dendrites in multiple sublaminae, including S1,3 and 5 (Figure 6C-E). Similar lamination patterns were revealed by labeling in a *Sst-cre* line (Figure 6F). Owing to the density of labeling, we were unable to determine whether individual ACs are multi-stratified or whether processes of individual C63 ACs are confined to a single sublamina.

### nGnG amacrines

As noted above four AC types expressed substantially lower levels of GABAergic and glycinergic markers than known GABAergic or glycinergic types (Figure 5A,B). We call these types nGnG-1-4. To characterize these types, we identified additional markers that distinguished them from each other and other ACs (Figure 7A). We used these markers to characterize them using immunohistochemistry and a set of transgenic lines.

**Figure 7.**
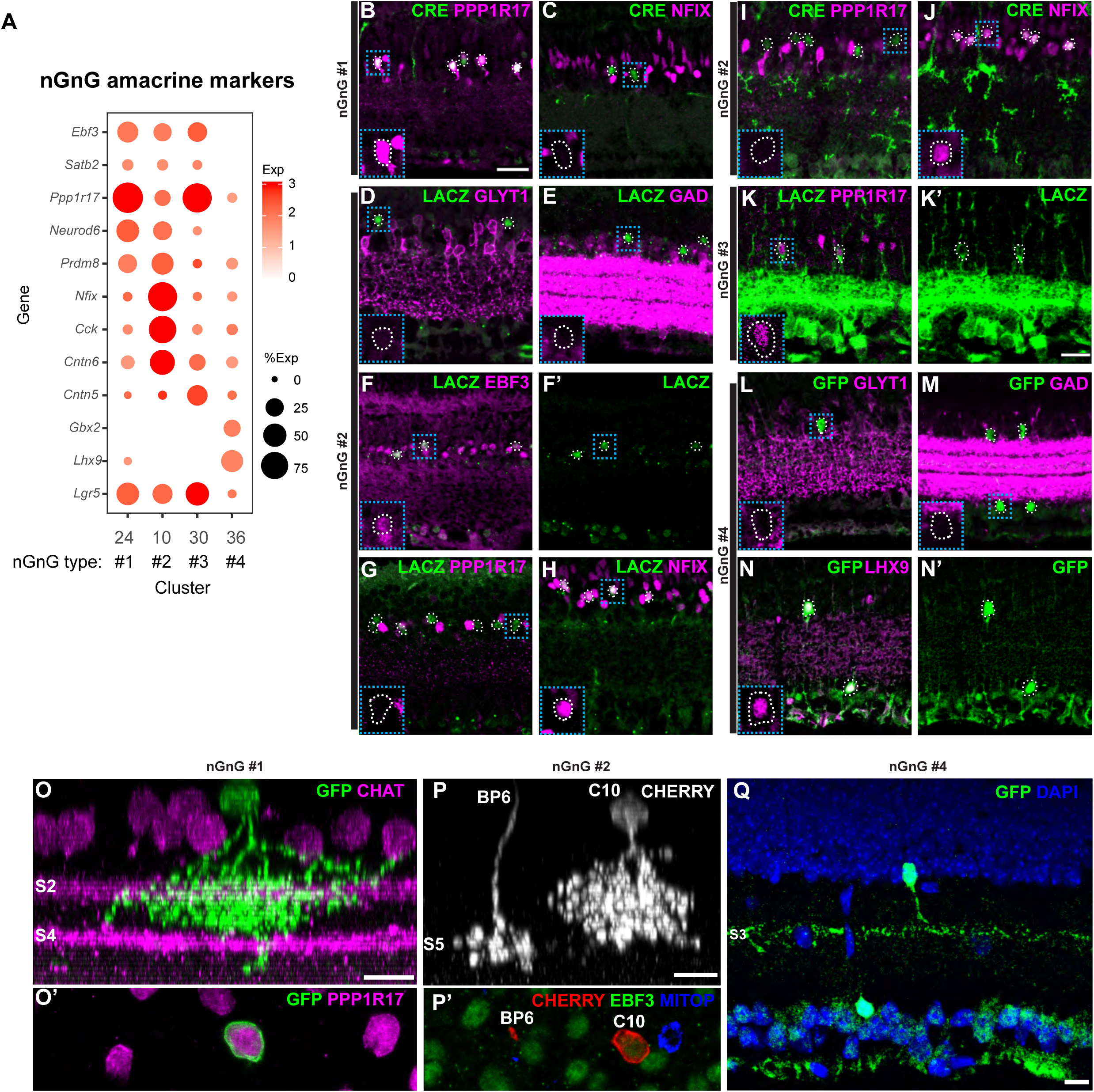
Molecular markers defining four types of nGnG ACs. A. Dot plot showing patterns of gene expression in four putative nGnG AC types (nGnG’s #1-4: C24, 10, 30 and 36). B-C. nGnG-1 ACs (C24) labeled by immunostaining *NeuroD6-cre* retinas with an antibodies against Cre recombinase together with either PPP1R17 or NFIX. They coexpressed PPP1R17 but were negative for NFIX. White dashed outlines indicate CRE-positive cells. Blue dashed square inset shows 2x magnification. Scale bar: 25µm. D-H. nGnG-2 ACs (C10) labeled in the *Cntn6-LacZ* line and detected by staining for LACZ were negative for GLYT1 (D) and GAD65/67 (E). They coexpressed EBF3 (F) and NFIX (H) but were negative for PPP1R17 (G). I-J. nGnG-2 ACs were also labeled using the *Cck-IRES-cre* and detected by staining for Cre recombinase. As when labeled by the *Cntn6-LacZ* line, most ACs labeled with this line were PPP1R17-negative and NFIX-positive. K. nGnG-3 ACs (C30) labeled in the *Cntn5-Lacz* line were detected by staining for LACZ and were PPP1R17-positive. L-N.nGnG-4 ACs (C36) labeled in the *Gbx2-CreER-GFP* line, were GLYT1-negative (L) and GAD65/67-negative (M) but LHX9-positive (N). O-Q.Dendritic lamination of nGnG ACs. O. nGnG-1 morphology revealed as PPP1R17 cells in the *Neurod6-Cre* line. nGnG-1 ACs were labeled by sparse viral infection using the AAV9-EF1a-BbTagBy brainbow virus. nGnG-1 ACs laminated S1-3, as demonstrated by CHAT staining (S2, S4). O’ shows a 90° rotated view of the labeled AC’s soma, confirming co-expression of PPP1R17. Scale bar: 10µm. P. nGnG-2 morphology revealed as EBF3-positive cells, *Mitop*-negative (a mouse line that labels nGnG-1) ACs in the *Cck-IRES-cre* line. nGnG-2 ACs were labeled by sparse viral infection using the AAV9-EF1a-BbCht brainbow virus. nGnG-2 ACs laminated in S1-4 (M,N), as demonstrated by its positioning relative to cone bipolar Type 6 axon terminals (S5), which are also labeled in this line. O’ shows a 90° rotated view of the labeled AC’s soma, confirming the labeled AC was EBF3-positive, Mitop-negative (detected by GFP staining). Scale bar: 10µm. Q. nGnG-4 lamination pattern revealed using the *Gbx2-CreER-GFP* line, GFP staining showed an AC that tightly laminated in S3. Scale bar: 10µm

nGnG-1 ACs (C24) expressed *Neurod6, Ebf3* and *Ppp1r17*, which we previously showed to define the nGnG ACs labeled in the *MitoP* mouse line (Kay et al., 2011, Macosko 2015). Consistent with previous results, it was labeled in a *Neurod6-Cre* line, as detected by immunostaining for Cre recombinase, co-expressed PPP1R17, and was negative for NFIX (Figure 7B, C).

nGnG-2 ACs (C10) expressed *Cntn6, Cck, Ebf3, Nfix* and *Prdm8*. We labeled them in *Cntn6-lacZ* and *Cck-IRES-Cre* lines and confirmed their nGnG status confirmed their nGnG status by immunostaining for GAD (using an antibody that recognizes both GAD65 and GAD67, encoded by Gad1 and Gad2) and GLYT1, observing they were negative for both and for PPP1R17, but positive for NFIX and EBF3 (Figure 7D-J).

nGnG-3 ACs (C30) expressed *Cntn5* and *Ppp1r17*. We labeled them in a *Cntn5-LacZ* line and showed that they co-expressed PPP1R17 (Figure 7K). The density of *Cntn5-LacZ* labeling in bipolar cells and RGCs precluded further examination of this cell type.

nGnG-4 (C36) expressed *Gbx2* and *Lhx9*. We labeled them in a *Gbx2-Creer-GFP* line and confirmed that both populations were GAD- and GLYT1-negative. Their somata were present in both the inner nuclear layer and ganglion cell layer. We additionally observed co-expression of LHX9 by immunostaining (Figure 7L-N).

Using a combination of transgenic mouse lines and immunohistochemical markers, we were able to discern the morphology of three of the nGnG types. nGnG-1 and 2 were narrow-field types (similar to glycinergic ACs, to which they were related; see above) with dendrites that arborize in S1-3 and S1-4, respectively (Figure 7O,P). nGnG-4 ACs had arbors tightly confined to S3 and appeared to be medium or wide field ACs, similar to GABAergic ACs, to which they were related) (Figure 7Q). The lamination pattern of these ACs is similar to that of CAII ACs (Vuong et al., 2015), which we believe correspond to C25, but they do not express detectable levels of TH or other catecholaminergic markers.

We noted that 3 of the nGnG types (nGnG1-3) expressed *Lgr5* (Figure 7A), a gene that has been studied intensively as a marker of stem and progenitor cells in multiple tissues, and a critical regulator their activity (Leung et al., 2018). It was recently reported that a set of Lgr5-positive ACs are able to reenter the cell cycle and generate new retinal neurons and glia (Chen et al., 2015). These might be nGnG ACs, consistent with their low level of canonical GABAergic and glycinergic AC markers. *Lgr5* is also expressed by two close transcriptional relatives of these nGnG ACs, C4 (SEGs) and C28, both of which are glycinergic; this expression is consistent with a report that many Lgr5+ ACs are glycinergic (Sukhdeo et al., 2014).

### Receptors for neurotransmitters and neuromodulators

ACs form synapses on BCs, RGCs and other ACs. We asked whether these cells express receptors for the transmitters and modulators that ACs might use. To this end, we used data from 46 RGC types (Tran et al., 2019) and 15 BC types (Shekhar et al., 2016) as well as the data from ACs generated in this study (Figures 8 and S2).

**Figure 8.**
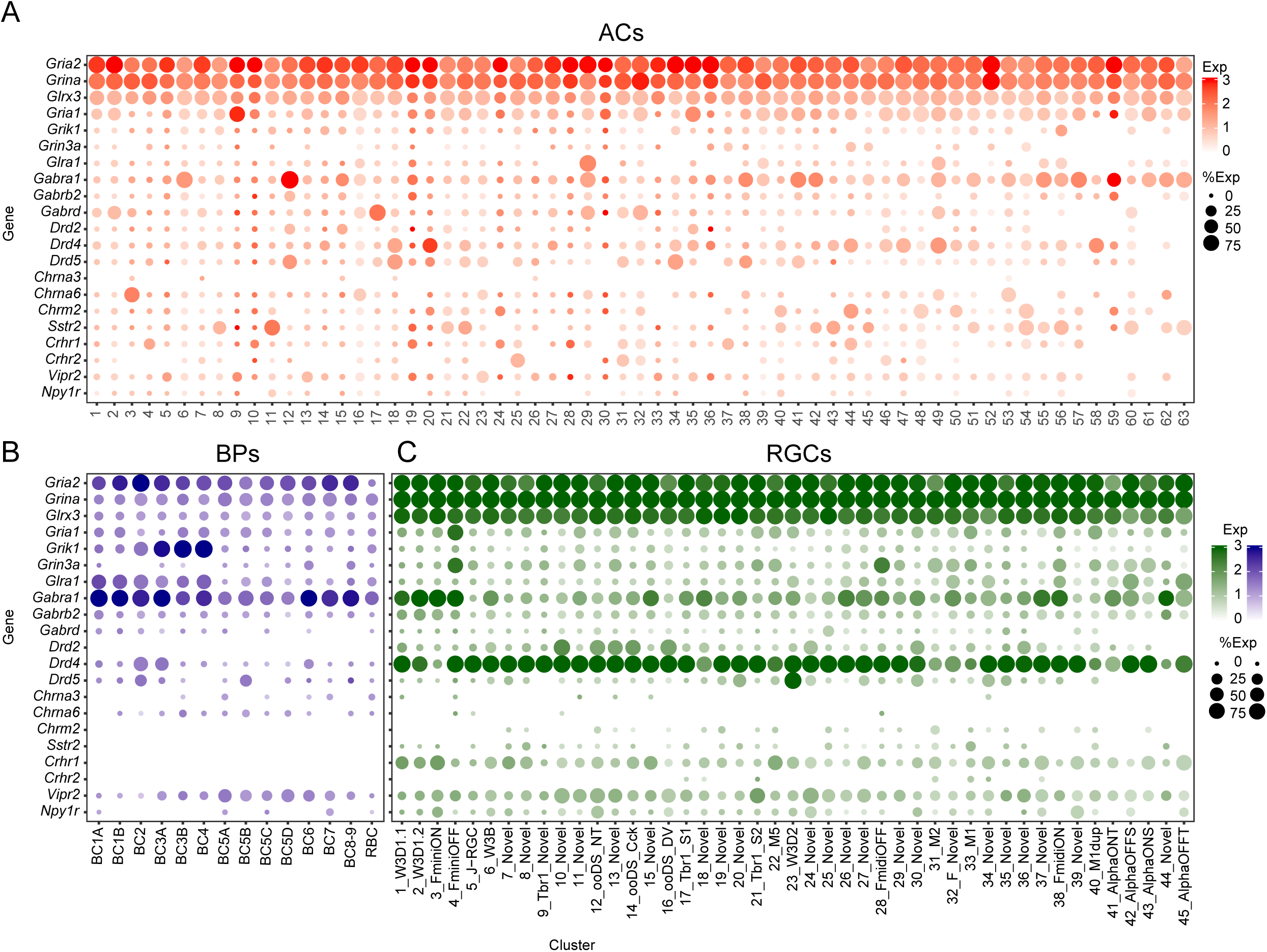
Expression of receptors for neurotransmitter and neuromodulator receptors in ACs, BCs, and RGCs. Examples of receptor expression in ACs (A, color in red), BCs (B, in dark blue) and RGCs (C, in dark green). Some receptors were broadly expressed in most cell classes, some were selectively to certain class, others were specific to certain types in certain classes.

In general, receptors were broadly expressed. All BC, AC and RGC types expressed at least some GABA, glycine and glutamate receptor subunits, consistent with the large number of GABAergic, glycinergic and glutamatergic neurons that form synapses in the inner plexiform layer. Perhaps more surprising, many acetylcholine and dopamine receptor subunits and many neuropeptide receptors were broadly expressed, even though only a few cell types use these transmitters or modulators, and their synapses are confined to a few sublaminae within the inner plexiform layer. For example, dopamine synthetic enzymes were expressed at high level in two amacrine clusters, but dopamine receptors were found in multiple types of ACs, BCs and RGCs. Likewise, although there is only a single cholinergic retinal cell type (SACs), both nicotinic and muscarinic acetylcholine receptors were expressed broadly.

Some receptor subunits were, however, selectively expressed by a small number of cell types. They include the glutamate receptor subunits *Gria1, Grik1* and *Grin3a*; glycine receptor Glyra1; GABA receptor subunits *Gabrd, Gabrb2* and G*abra1*; Dopamine receptors *Drd2, 4*, and *5*; cholinergic receptors *Chrna3, 6* and *Chrm2*; and neuropeptide receptors *Sstr2* and *Npy1r*. Interestingly, some adrenergic and serotonin receptors were also selectively expressed even though there is no evidence for the use of norepinephrine or serotonin as a retinal neurotransmitter in mice.

### Transcription factors defining transcriptionally related groups

The ability to order AC types by transcriptomic similarity (Figure 5A) led us to ask if we could identify transcription factors expressed by groups of related types (Figure 9A). We found a few transcription factors expressed by single AC types (*Sox2* [Whitney et al., 2014], *Mafb* and *Nfib*) and others expressed by small groups of related types (e.g., *Ebf3, Satb2* [Kay et al. 2011] and *Etv1*). Many transcription factors expressed by ACs were also expressed by subsets of RGCs (Tran et al., 2019). Of particular interest in this context were transcription factors selectively expressed by either GABAergic or glycinergic types. We focused on four such factors: *Meis2, Tcf4, Eomes* and *Neurod2*.

**Figure 9.**
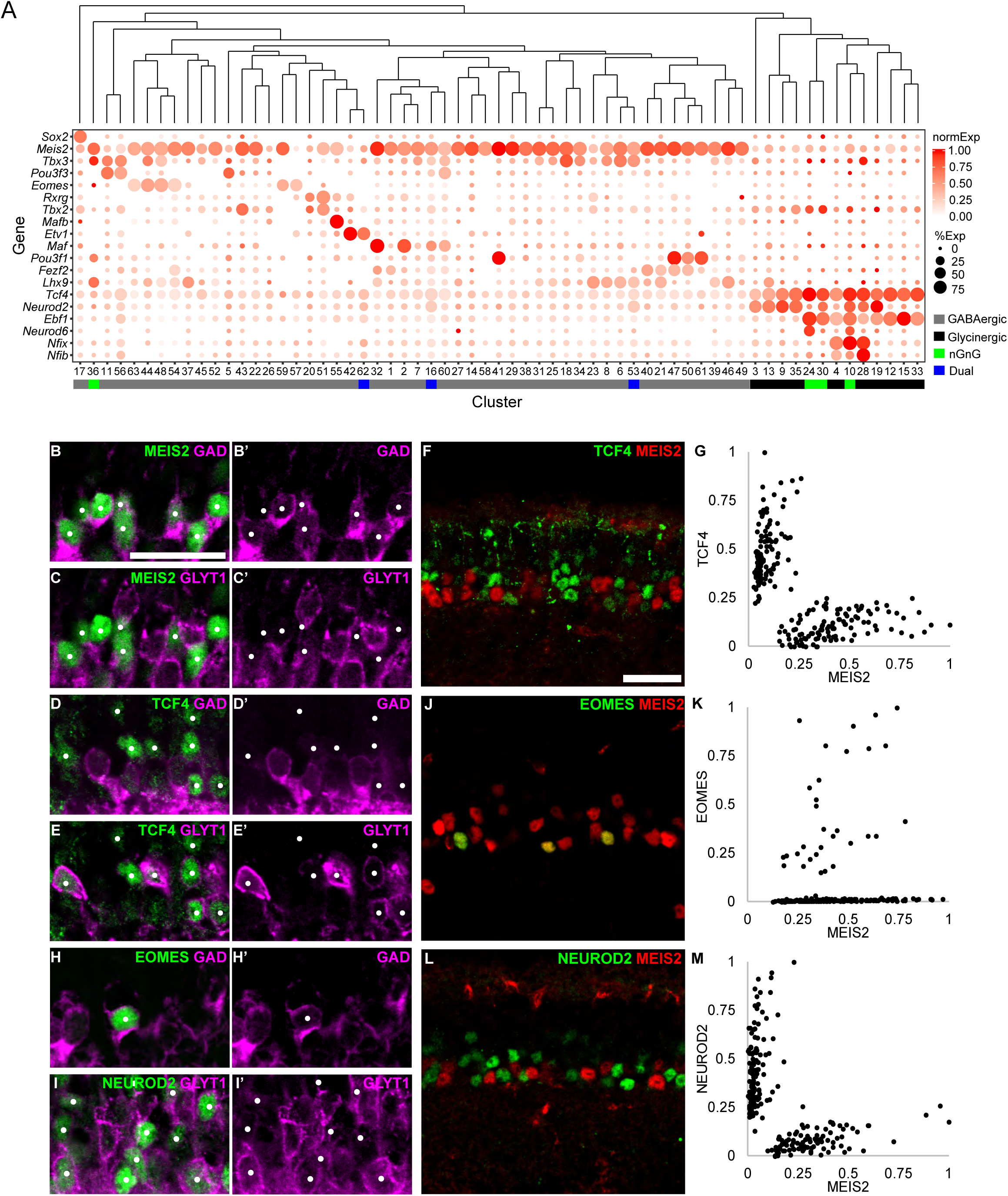
Transcription factors expressed by ACs. A. Expression of selected transcription factors by AC types. Dendrogram shows transcriptional relationships among AC types, generated by hierarchical clustering of average gene signatures (Euclidean distance metric, average linkage). Color indicates average normalized transcript level per cluster in expressing cells. B-C. Immunostaining shows that MEIS2-positive ACs (white dots) are GAD-positive (detected by an anti-GAD65/67 antibody) and GlyT1-negative in P21 mouse retina INL. F. MEIS2 and TCF4 mark mutually exclusive groups of ACs. G. Quantification of fluorescent intensity of MEIS2 (x-axis) and TCF4 (y-axis) positive cells in the INL at P21. Raw fluorescent intensity values were background subtracted and normalized to the maximum intesity value for each marker. (225 cells scored) H. EOMES-positive cells in the INL are GAD-positive. I. The majority of NEUROD2-positive cells in the INL are GLYT1-positive. J. EOMES-positive ACs are a subset of MEIS2-positive ACs. K. Quantification of fluorescent intensity of MEIS2 (x-axis) and EOMES (y-axis) positive cells in the INL at P21. (217 cells scored) L. NEUROD2-positive ACs are MEIS2-negative. M. Quantification of fluorescent intensity of MEIS2 (x-axis) and NEUROD2 (y-axis) positive cells in the INL at P21. (222 cells scored) Scale bars are 25µm.

*Meis2* was expressed at higher levels by most GABAergic types than by any glycinergic type (Figure 9A), a pattern supported by immunohistochemical analysis using the canonical GABAergic and glycinergic markers. Retinal sections were triple labeled with MEIS2, GAD65/67, and GLYT1 and fraction of co-labeling was calculated. MEIS2 was present in 78.1±2.3% mean ±SEM from ≥5 images from 2 animals) of GAD-positive ACs and 94.3±1.7% of MEIS2-positive ACs co-expressed GAD (332 cells scored). 2.0±0.6% of MEIS2-positive cells co-expressed GAD and GLYT1, suggesting they could be dual neurotransmitter ACs, and 3.2±1.0% were double-negative (Figure 9B,C). In contrast, *Tcf4* was expressed at higher levels by all glycinergic and nGnG-1-3 types than by any GABAergic type (Figure 9A). Immunostaining confirmed TCF4 expression in 83.0±4.2% of GLYT1-positive ACs and 71.7±3.8% of TCF4 were GLYT1-positive (417 cells scored) (Figure 9D,E). 25.0±3.4% of TCF4-positive cells were GLYT1-negative and GAD-negative, these likely represented nGnG ACs and 3.4±1.1% of TCF4-positive cells appeared to co-express GAD and GLYT1. Double-staining with anti-MEIS2 and anti-TCF4 and quantification of immunoreactivity levels confirmed that they were present in mutually exclusive AC subsets (Figure 9F,G).

*Eomes* (TBR2) and *NeuroD2* were expressed by restricted subsets of GABAergic and glycinergic AC types, respectively (Figure 9A). *Eomes*, previously studied as a marker for intrinsically photosensitive RGCs (Mao et al., 2014; Sweeney et al., 2014), was expressed at higher levels by six GABAergic types than by any glycinergic types, and all EOMES-positive ACs were GAD-positive (Figure 9H). As expected, all EOMES-positive ACs were also MEIS2-positive (Figure 9J,K). *NeuroD2* was expressed at higher levels by 6 glycinergic and 2 nGnG types than by any GABAergic types. The majority of NEUROD2 ACs expressed GLYT1, confirming a previous report demonstrating NEUROD2 expression in a subset of glycinergic ACs (Cherry et al. 2011) (Figure 9I). NEUROD2-positive ACs were mutually exclusive from MEIS2-positive ACs, suggesting the GLYT1-negative, NEUROD2-positive ACs are likely to be nGnG ACs (Figure 9L,M).

Together, these patterns suggest possible roles for MEIS2, TCF4, EOMES and NEUROD2 in establishing or maintaining the GABAergic or glycinergic phenotypes, or correlated features of the ACs that express them.

## DISCUSSION

### Mouse AC types

The heterogeneity of ACs has been recognized since the time of Cajal (1893) and documented by multiple methods (Diamond 2017). Minimally biased cell filling methods identified 29 AC types in rabbit (McNeil and Masland, 1998) and at least 25 in mouse (Badea and Nathans, 2004; Majumdar et al., 2009; Pang et al., 2012), and reconstruction from serial electron microscopic sections revealed 45 types in mouse retina (Helmstaedter et al. 2013). These studies were likely underpowered, in that they surveyed a few hundred cells (by light microscopy) or reconstructed a limited area (electron microscopy). Our survey, based on 32,523 cells, increases the number to 63 AC types (Table 1).

Although many AC types have been studied previously, and we have matched molecular to morphological features for others, the majority remain uncharacterized. Surprisingly, of the 10 most abundant types, 7 have not, to our knowledge, been subjects of previous studies (C1,2,5,7-9; Table 1). The markers we have identified may provide a starting point for investigations of these ACs.

Is the catalogue now complete? We may have missed AC types for at least three reasons. First, types comprising <0.1% of ACs might not have been detected, either because they were not collected or because our computational methods nominate clusters only when many cells share a transcriptional pattern. Second, we may have failed to collect some types for technical reasons – e.g., if they were particularly fragile. Third, some clusters could contain more than a single type. For example, our computational methods distinguish ON and OFF SACs as separate types in neonates (Peng et al., 2020) but their transcriptionally differences diminish with age, and they form a single cluster at P18-19. Thus, we view 63 as a lower limit to the number of AC types, but have no reason to expect that the true number greatly exceeds 63.

### Neurotransmitters and neuromodulators

ACs are largely inhibitory, using GABA and glycine as neurotransmitters (Masland 2012; Wassle et al., 2009; Diamond, 2017). Our results support this dogma, with 56 of the 63 AC types being either GABAergic (43) or glycinergic (13) based on expressing *Gad1/2* or *Glyt1*. In addition, however, we found 3 types that express both *Gad1/2* at levels found in most GABAergic ACs and *Glyt1* at levels found in some glycinergic types; these could use both transmitters. Although one of these types could be artifactual (composed to two ACs that were profiled together; see Results) it is unlikely for the others (see Methods). We also found 4 types that express low levels of both *Gad1/2* and *Glyt1*. These non-GABAergic non-glycinergic types, which we call nGnG1-4, include the type that we described previously (Kay et al., 2011; Macosko et al., 2015) as well as 3 additional types. We confirmed lack of detectable Gad and Glyt1 immunohistochemically, but physiological studies will be needed to determine whether low expression levels detected at the RNA level could nonetheless mediate GABAergic or glycinergic transmission. We found no evidence for the use of other small molecule neurotransmitters by these ACs, but all AC types, including nGnG ACs express neuropeptides.

In addition to GABA and glycine, some AC types have been shown to use glutamate, acetylcholine, dopamine, or NO, generally in addition to either GABA or glycine (Diamond, 2017). We identified AC types likely to use all of these transmitters, including a potentially novel glutamatergic type (VG1). The glutamatergic VG3 type is also glycinergic, as shown physiologically (Lee et al., 2016; Tien et al., 2016) whereas the VG1 type is likely to be GABAergic. We found no evidence that specific AC types synthesize other small molecule neurotransmitters, including serotonin, tyramine, histamine, epinephrine or norepinephrine. Some of these have been reported to be used by ACs in other species (Ghai et. al, 2009) but in the majority of cases, conclusions are based on uptake of or responsiveness to exogenous transmitter rather than production of endogenous transmitter (e.g., Fletcher and Wassle, 1997).

In summary, ACs use a remarkable array of neurotransmitters and neuromodulators, and most AC types are likely capable of releasing at least two such bioactive species, thereby enhancing the range of signals they can provide (Nusbaum et al., 2017). Finally, although most neurotransmitter and neuropeptide receptors were broadly expressed in ACs, BC, and RGCs, some were selectively expressed by a small number of neuronal types. Comparing their expression with that of neurotransmitter synthetic enzymes and neuropeptide genes in ACs can help to guide attempts to elucidate synaptic connectivity.

### Transcriptional relationships among AC types

Arranging AC types by transcriptional similarity (Figure 5,9) revealed several interesting relationships. First, the most fundamental (highest level) division is into GABAergic and glycinergic types. It is noteworthy that these two subclasses differ in additional ways: most GABAergic ACs have relatively broad dendritic arbors confined to one or a few sublaminae in the IPL, whereas most glycinergic ACs have narrow arbors that span multiple sublaminae (Wässle et al., 2009; Diamond, 2017). Thus, global comparison of genes differentially expressed by GABAergic and glycinergic types, such as the transcription factors discussed below, could reveal determinants of their contrasting morphologies as well as their transmitter choice.

Second, among GABAergic and glycinergic types, the most divergent are SACs and Aii ACs respectively. In fact, although the GABAergic nature of SACs is indisputable, they are no more closely related to other GABAergic than to glycinergic types. SACs are more closely related to RGCs than other ACs (Macosko et al. 2015), are among the first-born during embryogenesis (Voinescu et al., 2011), and play an organizing role in patterning the sublamination of the IPL (Peng et al., 2017; Duan et al., 2018). Aii, the most divergent of glycinergic ACs, play a unique role in providing the principle route through which input from rods is delivered to retinal ganglion cells (Demb and Singer, 2012). We speculate that SACs and Aii ACs may be among the most evolutionarily ancient of AC types.

Third, some AC types with similar transmitter profiles are close transcriptional relatives, but this is not always the case. For example, three of the nGnG types are close relatives of each other but one type is distant. nGnG-1 is also closely related to the glycinergic SEG AC, consistent with our previous demonstration that a postmitotic fate choice diversifies these two (Kay et al., 2011). Some peptide-expressing clusters are closely related – for example, two of the VIP-positive types and two of the CCK-positive types. In contrast, shared expression of other neurotransmitters or neuromodulators is not reflected in overall transcriptional similarity. Thus, two other CCK-expressing types are distant relatives of each other and the two putative glutamatergic types (VG1 and VG3) are not close relatives.

Fourth, several transcription factors, including *Meis2, Tcf4, Ebf1, Neurod2* and *Eomes* (*Tbr2*) are expressed by groups of closely related GABAergic (*Meis2, Eomes*) or glycinergic (*Tcf4, Ebf1, Neurod2*) AC types. Association of some of these genes with GABAergic (Meis2: Bumsted-O’Brien et al., 2007) or glycinergic (NeuroD2: Cherry et al., 2011) ACs has been noted previously but comprehensive analysis of their expression has not been reported. Of these, *Meis2* and *Tcf4* are of particular interest, since they are expressed by most GABAergic and all Glycinergic ACs, respectively. Both transcription factors play critical developmental roles in multiple cell types, both within and outside of the nervous system (*Tcf4*: Forrest et al., 2014; *Meis2*: Schulte and Geerts, 2019) but their roles in retinal development have not been reported to date.

The patterns we documented raise the tantalizing possibility that selectively expressed transcription factors may plays roles in specifying AC types or subclasses, or determining features they share. It will be possible to test these possibilities using genetic methods that have been applied successfully to other transcriptional regulators in other retinal cell types.

### The mouse retinal cell atlas

We previously analyzed all retinal cell classes other than amacrines, documenting the existence of 65 neuronal types and 6 non-neuronal types (astrocytes, endothelial cells, fibroblasts, microglia, Müller glia and pericytes) (Macosko et al., 2015; Shekhar et al., 2016; Tran et al., 2019). To these we now add 63 AC types. Thus, the mouse retinal atlas stands at 134 cell types (Table 2).

**Table 2.** The mouse retinal cell atlas. Number of types per class estimated in 2010 (Wässle et al., 2009; Masland, 2012) and current estimate based on data in this study, Macasko et al. (2015), Shekhar et al. (2016) and Tran et al. (2019).

| **Class** | **# Type in 2010** | **# Type in 2020** |
| --- | --- | --- |
| **PR** | 3 | 3 |
| **HC** | 1 | 1 |
| **BC** | 12 | 15 |
| **RGC** | 20 | 46 |
| **AC** | 29 | 63 |
| **Total** | 65 | 128 |

The existence of 6 major classes of retinal cells (5 neuronal plus Müller glia) was clear nearly 130 years ago (Cajal, 1893). Over the subsequent 120 years, classical studies in numerous labs and of numerous species defined many types within classes, but the low-throughput nature of the available methods prevented generation of a comprehensive inventory. In an influential and authoritative review published less than a decade ago, the number of retinal cell types was estimated to be under 70 (Masland, 2012) (Table 2). Since that time, the estimated number has nearly doubled, largely as a result of applying newly developed high-through ultrastructural, physiological and, above all, transcriptomic methods. Although it is too soon to declare victory, we believe that the current number is close to accurate.

## Acknowledgments

We thank Drs. Thomas Bourgeron (Institut Pasteur), Isabelle Cloez-Tayarani (Institut Pasteur), and James YH Li (University of Connecticut) for providing some mouse tissue used in this study; and Drs. Julia Kaltschmidt (Stanford University), Louis Reichardt (Simons Foundation), Chinfei Chen (Harvard Medical school), Lisa Goodrich (Harvard Medical school) for providing mice. This work was supported by the NIH (NS029169, MH105960, K99EY029360), and Human Frontiers Science Program).

## Extended Data

**Figure 6-1.**
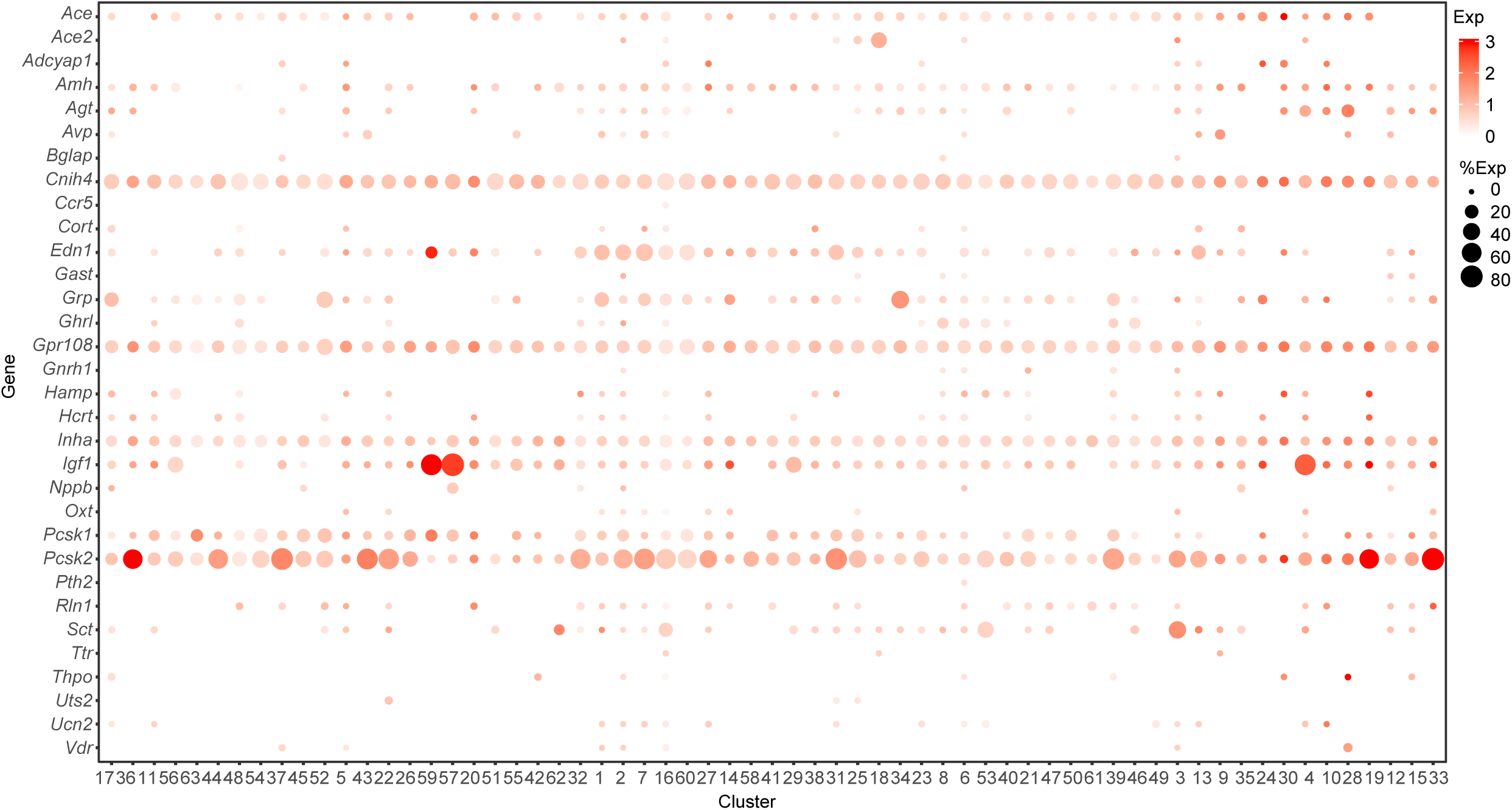
Expression of neuropeptides in ACs.

**Figure 8-1.**
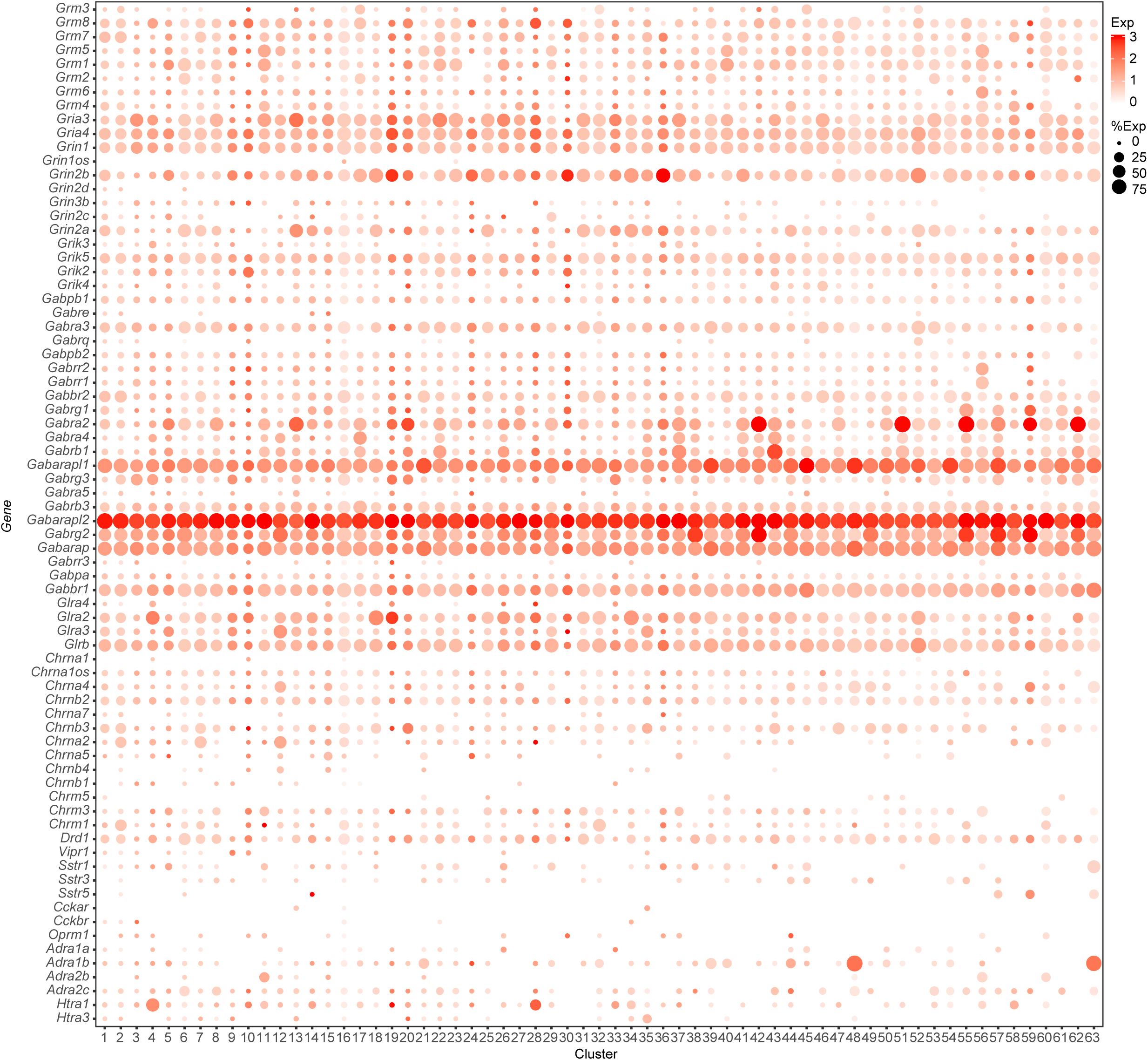
Expression of neurotransmitter and neuropeptide receptors in ACs.

